# The relationship between frequency content and representational dynamics in the decoding of neurophysiological data

**DOI:** 10.1101/2022.02.07.479399

**Authors:** Cameron Higgins, Mats W.J. van Es, Andrew Quinn, Diego Vidaurre, Mark Woolrich

**Affiliations:** Oxford Centre for Human Brain Activity, Wellcome Centre for Integrative Neuroimaging, Department of Psychiatry, University of Oxford, Oxford, UK; Center of Functionally Integrative Neuroscience, Department of Clinical Medicine, Aarhus University, Aarhus, Denmark

## Abstract

Decoding of high temporal resolution, stimulus-evoked neurophysiological data is increasingly used to test theories about how the brain processes information. However, a fundamental relationship between the frequency spectra of the neural signal and the subsequent decoding accuracy timecourse is not widely recognised. We show that, in commonly used instantaneous signal decoding paradigms, each sinusoidal component of the evoked response is translated to double its original frequency in the subsequent decoding accuracy timecourses. We therefore recommend, where researchers use instantaneous signal decoding paradigms, that more aggressive low pass filtering is applied with a cut-off at one quarter of the sampling rate, to eliminate representational alias artefacts. However, this does not negate the accompanying interpretational challenges. We show that these can be resolved by decoding paradigms that utilise both a signal’s instantaneous magnitude and its local gradient information as features for decoding. On a publicly available MEG dataset, this results in decoding accuracy metrics that are higher, more stable over time, and free of the technical and interpretational challenges previously characterised. We anticipate that a broader awareness of these fundamental relationships will enable stronger interpretations of decoding results by linking them more clearly to the underlying signal characteristics that drive them.

**Highlights:**

- We investigate different decoding paradigms applied to epoched data and characterise the information content available to each over time.
- Under commonly used instantaneous signal decoding paradigms, sinusoidal components of the evoked response are translated to double their original frequency in decoding accuracy metrics, presenting technical and interpretational challenges.
- When instantaneous signal decoding is used, we recommend using low pass filters with a cut-off less than one quarter of the sampling rate to eliminate spurious representational alias artefacts.
- The interpretational issues associated with instantaneous signal decoding can be resolved with alternative paradigms such as complex spectrum decoding.
- We show that complex spectrum decoding results in decoding accuracy metrics that are higher, more stable over time, and free of representational aliasing.

## INTRODUCTION

The field of representational dynamics uses temporal patterns in decoding accuracy timecourses to test hypotheses about how the brain processes information (T. Carlson et al., 2013; Cichy et al., 2014; Kietzmann et al., 2019; King & Dehaene, 2014). By decoding different experimental stimuli from recorded brain activity at high temporal resolution, researchers use information theoretic measures to quantify what features of a stimulus are explicitly represented in neural data as a function of time from stimulus onset (T. A. Carlson et al., 2011; Cichy et al., 2016; Ince et al., 2017). An emerging question in neuroscience is how these representational dynamics relate to the brain’s underlying neurophysiology (Gross et al., 2013; Jafarpoura et al., 2013; Kriegeskorte & Kievit, 2013; Schyns et al., 2011). Such analyses seek to go beyond merely answering *what* is represented in recorded brain activity, by also characterising the neural mechanisms explaining *how* that information is represented (Higgins, Vidaurre, et al., 2021; Kikumoto & Mayr, 2018; Valentin et al., 2020; van de Nieuwenhuijzen et al., 2013; Zhan et al., 2019).

This commonly involves a decoding paradigm we will refer to as *instantaneous signal decoding*, where classifiers are trained and tested on the raw broadband signal recorded over all sensors at each timepoint following a stimulus (T. Carlson et al., 2013; Cichy & Pantazis, 2017; Grootswagers et al., 2017), and the representational dynamics interpreted (often with reference to activity in canonical frequency bands). This can be used for example to study the phase-locking of information content to canonical oscillations (Kerrén et al., 2018; Kunz et al., 2019; van Es et al., 2020), the dynamics of memory (Higgins, Liu, et al., 2021; LaRocque et al., 2013; Wolff et al., 2015), or the direction of information flow (Cichy et al., 2014; Dijkstra et al., 2020; Goddard et al., 2016; Linde-Domingo et al., 2019). A closely related paradigm, we will refer to as *narrowband signal decoding*, applies the same procedure after filtering the data into a narrowband of interest. This explicitly links observed patterns with canonical frequency bands (Samaha et al., 2016; Xie et al., 2020).

Unfortunately, however, the fundamental relationship between the frequency content of the stimulus evoked signal and the inferred information content is not widely recognised. Whilst many decoding approaches aim to be agnostic about the specific data characteristics over time that drive their results, there is a considerable risk of misinterpretation when this relationship is not considered. In this paper we draw attention to this relationship, highlighting that the spectral content of the evoked response is translated to double its original frequency in associated decoding accuracy metrics when using the *instantaneous signal decoding* or *narrowband signal decoding* paradigms most typically used in the literature (T. Carlson et al., 2013; Cichy & Pantazis, 2017; Grootswagers et al., 2017). From this, we identify two problems: the first is the presence of artefacts due to representational aliasing; the second is the broader challenge of how we should interpret information theoretic metrics that systematically oscillate at double the frequency of the evoked response spectrum.

We argue that these problems arise from a narrow focus on information content in the instantaneous signal at a single moment in time, which ignores information stored in the signal’s gradient or higher moments. Conceptually, this is analogous to analysing a simple pendulum by measuring only its displacement at a single instant in time – not its velocity or acceleration, which would together fully define the dynamic system. As illustrated in figure 1, such a narrow focus only on the pendulum’s displacement leads to inferred information content that peaks at the pendulum’s extrema (i.e. the peaks and troughs of the oscillation); taking a broader view of the information contained in both the displacement and velocity leads to a measure of information content that is stable over time.

**Figure 1:**
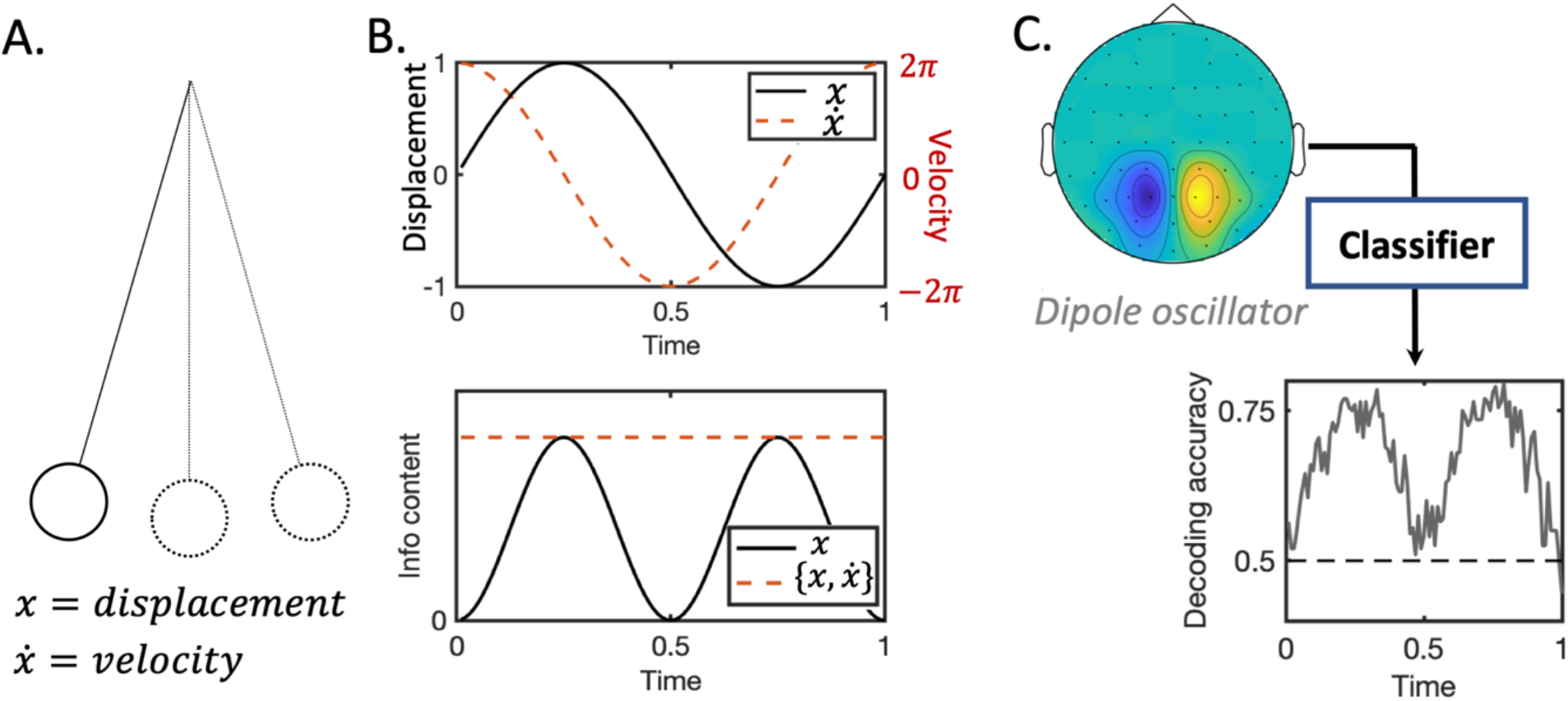
a pendulum analogy for decoding oscillatory neural signals. **A**. Commonly used *instantaneous signal decoding* pipelines can only offer a partial view of the brain’s representational dynamics, as they only use the instantaneous data values and cannot detect information stored in the gradient or higher moments of the dynamic signal trajectory. When dealing with a dynamical system such as the brain, this is like trying to predict the behaviour of a pendulum given only its displacement at a single instant in time – not its velocity or momentum, which would fully characterise the dynamic system. **B**. Suppose we wish to classify if a pendulum is moving or stationary given noisy estimates of its displacement and velocity over time. The information content associated with only the displacement readout peaks at the pendulum’s extrema and drops to zero in between. Including the velocity information instead achieves stable information content over time. **C**. Evoked neural data with strong oscillatory components behaves in the same way as the pendulum. When researchers apply *instantaneous signal decoding* to such data, classifiers should perform well at the peaks and troughs of sinusoidal components in the evoked response, and poorly in between. This problem would be overcome if the local gradient information was included as features for classification, resulting in information content metrics that are stable over time.

We extend the same logic to the dynamic trajectory of neural activity evoked by a stimulus. This motivates a third decoding paradigm that we refer to as *complex spectrum decoding*, which is one way of including such temporal gradient information. Returning to our example above, if we applied a Fourier decomposition to the pendulum’s displacement over time, we would obtain a single complex frequency component with a real part (tracking the displacement) and an imaginary part (tracking the velocity). This concept generalises to neural activity, where we would expect more complex Fourier dynamics played out simultaneously over multiple frequency bands and spatial channels. When this complex spectrum information is included as features to a classifier, we show that this results in inferred representational dynamic patterns that have higher accuracy, are more stable over time, and which we believe to provide a better characterisation of the brain’s representational architecture.

## RESULTS

### 1. How the spectrum of the evoked response determines the signal information content

We first ask: what is the fundamental relationship between frequency-specific features of the stimulus-evoked response and the resulting timecourse of decoding accuracy? We address this question using a generative modelling approach, where we model the neural data recorded on individual trials as a Fourier series with bandlimited Gaussian noise. From a probabilistic modelling perspective, the *mutual information* is the theoretical quantity analogous to decoding accuracy that we can then derive. This allows us to characterise how the information content of a signal varies as a function of time and frequency.

#### 1.1. Generative model of stimulus evoked responses

We wish to model epoched electrophysiological data recorded from *P* channels under two different experimental conditions. Let us denote by *x*_*n,t*_ the [*P* × 1] vector of data recorded at time *t* ∈ {1,2, … *T*} on trial *n* ∈ {1,2, … *N*}, where *y*_*n*_ ∈ {1, −1} denotes the experimental condition for that trial. We model *x*_*n,t*_ as comprising a condition-independent evoked response term *μ*_*t*_ of dimension [*P* × 1], and residual terms that are decomposed into a sum of [*P* × 1] Fourier components *z*_*n,t,ω*_:

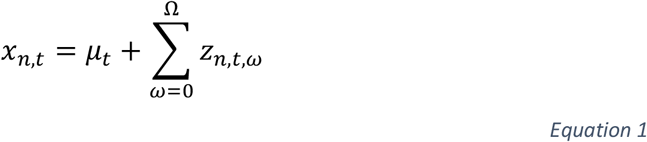

We henceforth refer to *x*_*n,t*_ as the ***‘broadband signal’***, and the multiple *z*_*n,t,ω*_ terms as the ***‘narrowband signals’***. If we assume each narrowband signal *z*_*n,t,ω*_ has a multivariate Gaussian distribution with mean conditioned on the stimulus (see Methods for full details), we obtain the following expression for the distribution of the broadband signal:

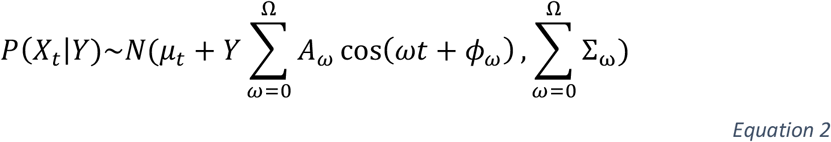

Each *A*_*ω*_ term is a diagonal [*P* × *P*] matrix, where the *i*th diagonal entry, denoted by *a*_*ω*,i_, reflects the magnitude of the component at frequency *ω* on channel *i*. Both *ω* and *t* are scalar indices reflecting the frequency and time respectively; *ϕ*_*ω*_ is a [*P* × 1] vector, each entry of which contains the phase offset of the oscillation at frequency *ω* across the *P* channels. Finally, we model induced effects (i.e. narrowband power that is not phase aligned to the stimulus) independently in each frequency band, where Σ_*ω*_ is the [*P* × *P*] covariance matrix modelling the spatial variance and correlations expressed at frequency band *ω*. Note that this corresponds to an assumption that only the evoked response, not the induced response, differs over the two conditions – this is a simplifying assumption that we later relax in Results Section 1.4.

We can now characterise the mutual information between the broadband signal *X*_*t*_ or its constituent narrowband signals *Z*_*t,ω*_and the class labels *Y*.

#### 1.2. Information content available to narrowband signal decoding

We wish to explore how the spectrum of the evoked response determines the representational dynamics inferred from the decoding paradigms that are most typically used in the literature (T. Carlson et al., 2013; Cichy & Pantazis, 2017; Grootswagers et al., 2017). We start by considering instantaneous decoding of narrowband signals *Z*_*t,ω*_, which we refer to as *narrowband signal decoding*.

Given a probabilistic model, we can calculate the *mutual information I*(*Z*_*t,ω*_, *Y*), which expresses the amount of information shared between the signal and the condition label time courses. This measure of information content in the signal that pertains to the condition labels can be thought of as a surrogate measure of decoding accuracy were one to do *narrowband signal decoding*. Starting with a single Fourier component of the evoked response at frequency *ω*, the mutual information is itself a sinusoidal function that has been translated to double the original frequency, 2*ω*:

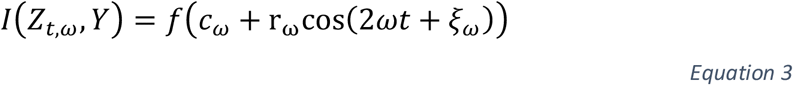

Where f is a monotonic, concave function (see Appendix B for proof and Figure A1 for plot of the function); and *c*_*ω*_, r_*ω*_ and *ξ*_*ω*_ are all scalar values that are constant with respect to time (see Appendix C for their exact values, and Appendix A and C for proof of the above result). The intuition for this is based on what was discussed in Figure 1: if *Z*_*t,ω*_ were the displacement of a pendulum oscillating at frequency *ω*, a decoder will perform best at the peaks and troughs of that oscillation and poorly in between these points.

We illustrate this relationship in example 1 (Figure 2), where we simulate an evoked response under two conditions. Suppose that one condition (in blue) contains information content at 10Hz across both channels, and the second condition (in black) does not. The information content associated with this narrowband component is itself a sinusoidal function oscillating at 20Hz.

**Figure 2:**
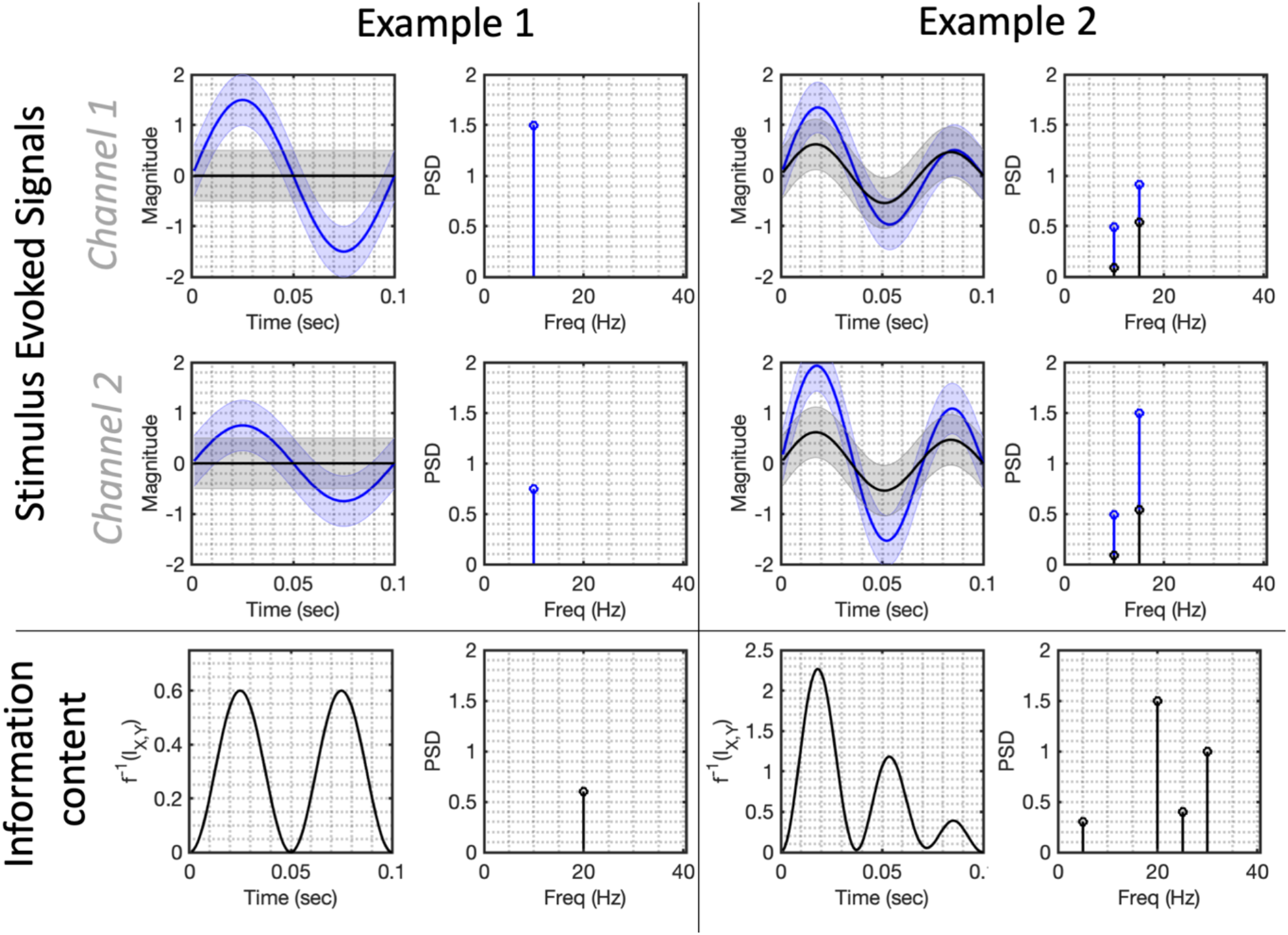
How the stimulus evoked spectrum determines the spectrum of information content when *instantaneous signal decoding* is used. In example 1 on the left, we simulate two conditions across two channels, with the upper panel showing the two conditions’ trial-averaged evoked responses (one in blue and one in black). The first condition evokes a phase-locked 10Hz oscillation on both channels, the second condition is a null condition in which there is no evoked response. The information content (i.e. the mutual information *I*(*X*_*t*_, *Y*) between the broadband data *X*_*t*_ and the stimulus labels *Y*) is plotted in the lower panel, and can be thought of as a surrogate measure of decoding accuracy were one to do *instantaneous signal decoding*. Although the only oscillation in the evoked response is at 10Hz, the information content is a 20Hz sinusoidal signal, reaching maxima at each peak and trough of the evoked response. In example 2 on the right, we simulate a signal comprised of 2 Fourier components at 10Hz and 15Hz in both conditions, with slightly different amplitudes. The associated information content is a signal with three distinct peaks, with Fourier components at 20Hz and 30Hz and additional harmonics at 5Hz and 25Hz.

#### 1.3. Information content available to instantaneous signal decoding

Realistic neural signals are not expressed in a single component frequency across all spatial areas, but are rather comprised of a number of spatially distinct components at multiple frequencies. How then does the entire frequency spectrum of the evoked response determine the frequency spectrum of the associated information content? This equates to the paradigm of *instantaneous signal decoding* that is most widely performed in the literature (T. Carlson et al., 2013; Cichy & Pantazis, 2017; Grootswagers et al., 2017). For the broadband signal *X*_*t*_ given in our model, the information content is given by:

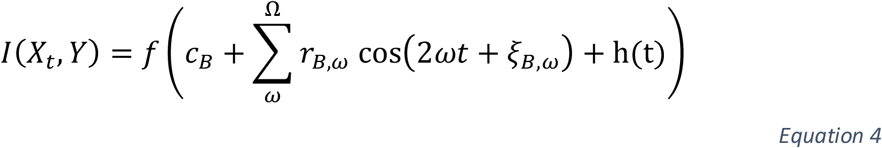

Where *c*_*B*_, *r*_*B,ω*_ and *ξ*_*B,ω*_ are scalar values that are constant over time, and h(t) refers to additional sinusoidal harmonics distributed across the frequency spectrum between zero and 2Ω (see Appendix D for their exact values along with proof of this result).

Importantly, if the highest frequency component of the evoked response on any channel is Ω, it follows that the highest frequency in the associated information spectrum will be 2Ω. We illustrate this point with example 2 in Figure 2; for simplicity we simulate an evoked response comprising just 2 spectral components under each condition at 10Hz and 15Hz; the associated information content displays multiple peaks over time, represented in its Fourier spectrum by frequency components distributed between 0Hz and 30Hz. As we will explore further in Results Section 2.1, this also means that commonly used anti-aliasing filters are insufficient to stop representational aliasing, i.e. alias artefacts in the inferred information content dynamics.

#### 1.4. Modelling induced effects

It is important to consider the degree to which these findings are specific to our chosen modelling assumptions. We have specifically limited our discussion to that of evoked effects by assuming the noise distribution was invariant over conditions. In the frequency domain, this means that we have limited our analysis to the part of the signal that is phase aligned to stimulus onset. When we introduce condition-specific induced effects – i.e. to model the case where one condition induces an increase in bandlimited power that has random phase alignment with the stimulus onset – we can no longer derive an exact analytic expression for the mutual information; however, we can derive an upper bound on the information content. This upper bound is a function of components at the same frequencies specified in equations 3 (for the narrowband case; see Appendix F and G for proof) and 5 (for the instantaneous signal case; see Appendix H for proof). This result is not mathematically trivial, but may nonetheless be intuitive to some readers on the basis that the information content of a signal containing both evoked and induced effects must be less than the combined information content of each of those effects assessed independently; and the information content of induced effects assessed independently is constant with respect to time (owing to the uniform phase distribution that defines induced effects). Thus, we are able to generalise our findings to the case where induced effects are present.

### 2. Technical and Interpretational issues raised

The relationship we have characterised above between the stimulus-evoked signal spectrum and the spectrum of the information content raises several issues with commonly used *instantaneous signal decoding* pipelines. On a technical level, there is a risk of high frequency artefacts which we refer to as representational aliasing. On a broader level, this raises questions about how certain features of decoding accuracy timecourses should be interpreted.

#### 2.1. Representational Aliasing

The Nyquist frequency defines the highest frequency component that can be correctly resolved from data that has been digitally sampled at a specified sampling rate. It is standard practice to apply a low pass anti-aliasing filter prior to sampling which ensures no *signal* components are above the Nyquist frequency and that all signal components can therefore be correctly resolved. However, this only applies to the signal components, not their associated information spectrum, which we have shown contains spectral contents at double the highest frequency of the signal spectrum.

It follows that representational aliasing artefacts will be present in *instantaneous signal decoding* accuracy metric unless the following condition is met:

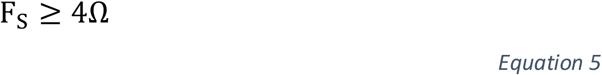

Where Ω is the highest frequency component of the evoked response and *F*_*s*_ is the sampling rate. Thus, *instantaneous signal decoding* pipelines need to use low pass filters with cut-off no higher than one quarter of the sampling rate – before training classifiers – in order to eliminate representational aliasing effects. Figure 3 illustrates this graphically.

**Figure 3:**
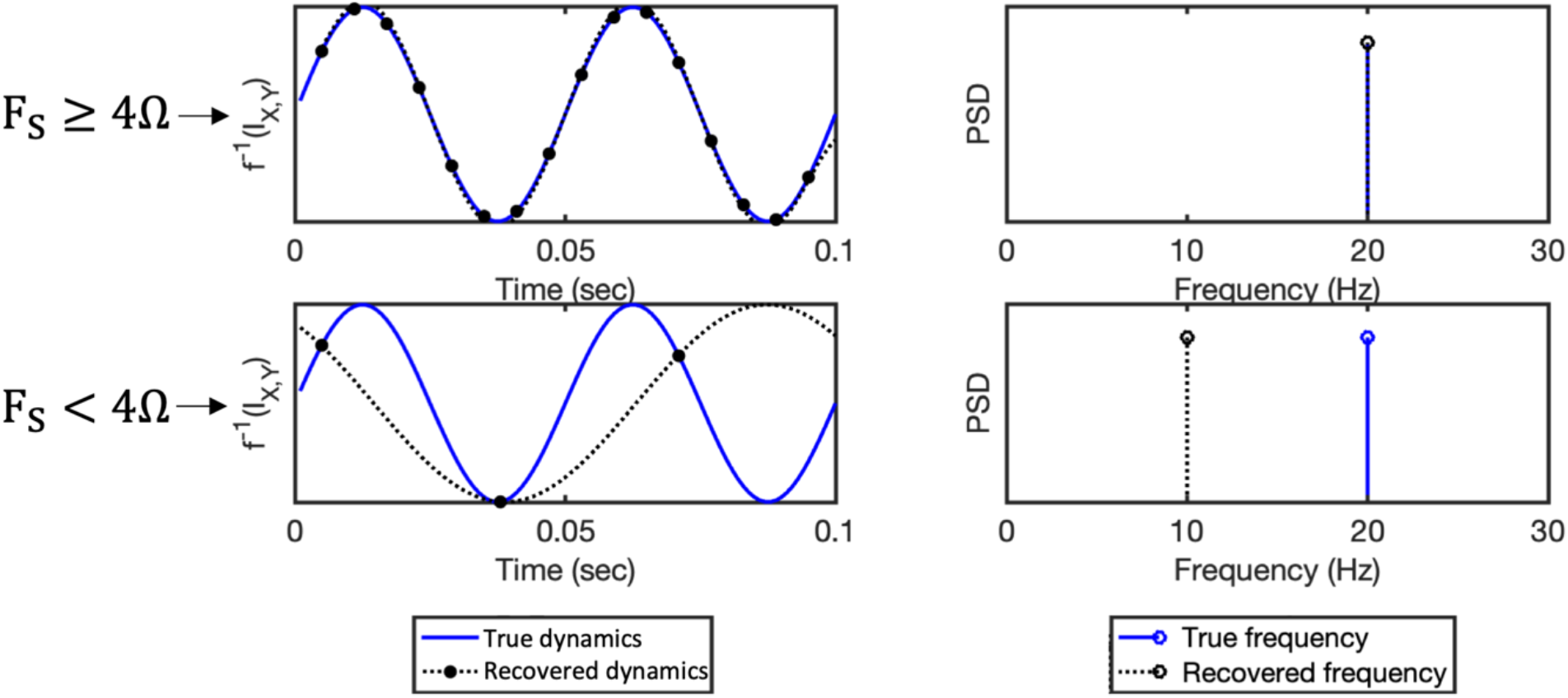
Demonstration of representational aliasing associated with *instantaneous signal decoding*. Consider the information content associated with example 1 from Figure 2, where one condition was associated with a stimulus-evoked component at 10Hz. As Figure 2 shows, the single oscillatory mode at 10 *Hz* is associated with a true information content that oscillates at a frequency of 20 Hz (plotted in blue above). In the first case on the top row, given a sampling rate of 160 *Hz* > 4 Ω, the recovered representational dynamics (plotted with black dashed line showing sinusoidal interpolation between the discrete samples) match the true frequency. In the second case however, given an inadequate sampling rate of 30 *Hz* < 4 Ω, the recovered dynamics are subject to representational aliasing, resulting in spurious dynamics at 10Hz rather than the true 20Hz pattern.

#### 2.2. How should we interpret oscillatory information content?

The oscillatory nature of information content associated with sinusoidal components of the evoked response is, we argue, interpretationally problematic. Features resembling successive peaks in classification accuracy are quite commonly reported in the literature (T. Carlson et al., 2013; Gennari et al., 2021; Hogendoorn & Burkitt, 2018; Kurth-Nelson et al., 2015; Mohsenzadeh et al., 2018; Robinson et al., 2020), where they have occasionally been interpreted as evidence for complex underlying phenomena – for example, the activation of distinct feedforward vs feedback connections, or discrete and distinct stages of cognitive processing. As we have shown in Figure 2, successive peaks arise naturally from an evoked response containing sinusoidal components. We argue that a simpler explanation for their common appearance in the literature could merely be that the typical evoked response is characterised by a succession of peaks and troughs (e.g. the N70, P100 and N175) that resemble a transient sinusoidal waveform.

We believe a fuller picture of information content should include the information stored in the dynamic gradient of the signal that is not available using *instantaneous signal decoding* pipelines. In Results Section 3 we explore a third paradigm that includes such information, and show that this results in narrowband information content that is stable over time. However, as these methods will not always be practical for reasons given in the discussion, we would more generally argue that representational dynamics obtained using *instantaneous signal decoding* and representing the ‘double peak’ feature shown in Figure 2 (and widely characterised in the literature) should first be assumed to correspond merely to peaks and troughs of an evoked sinusoidal signal, rather than more complex cognitive phenomena.

### 3. Obtaining measures of sinusoidal information content that are stable over time

We contend that the profile of information content obtained by *instantaneous signal decoding* is potentially misleading, as it suggests the brain’s representational dynamics are much faster than the actual evoked spectrum from which they are derived. Whilst *instantaneous signal decoding* pipelines are the most popular way to apply decoding to neural data at high temporal resolution, alternative methods exist that overcome these limitations. We focus our attention on Fourier analysis (for continuity with our modelling approach and because of these methods are well-established in neural data analysis), but emphasise these benefits are not specific to Fourier analysis per se – rather, they arise whenever methods include information in a dynamic signal’s higher moments (e.g. its gradient and rate of change) as features for classification.

#### 3.1. Complex Spectrum Decoding

We previously characterised the information content between stimulus labels *Y* and the narrowband Fourier series components *Z*_*t,ω*_. These narrowband components do not in fact include all the information that is returned by a Fourier signal decomposition; they reflect only the real component of a complex number representation. The imaginary components of these narrowband components reflect the instantaneous gradient information of each narrowband signal; we here characterise the information content associated with the full complex signal representation of each narrowband component, analogous to the decoding accuracy that would be obtained when both the narrowband signal and its local gradient are used as features for classification.

##### 3.1.1. Real and complex components of a Fourier decomposition

Fourier decompositions provide a complex representation of the underlying signal that includes both a real signal component and an orthogonal imaginary component, which we omitted from our model outline in Results Section 1 for simplicity. Including this complex-valued information, the same model can equivalently be written:

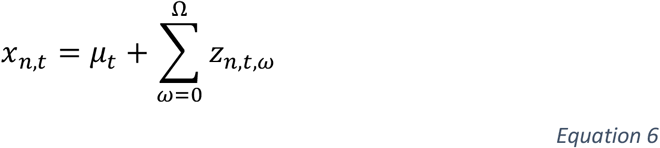

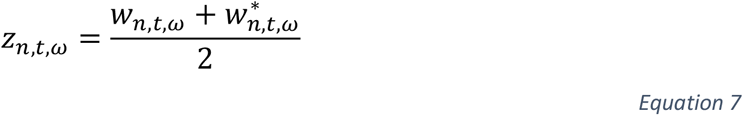

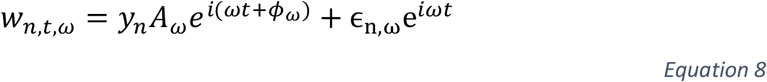

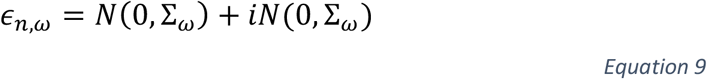

Where 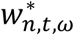 denotes the complex conjugate of *w*_*n,t,ω*_. This is exactly equivalent to the model of Equation 2, however it includes the complex spectral representation *w*_*n,t,ω*_ of each narrowband Fourier series component. It includes a condition-dependent evoked term 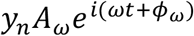 (i.e. the component of the response that is phase-locked to the stimulus), and a condition-independent residual term (i.e. the residual component with randomly drawn phase and amplitude on each trial; note that the values for the phase and amplitude are respectively the angle and magnitude of the complex valued *ϵ*_*n,ω*_ converted to polar coordinates).

##### 3.1.2. Information content available to complex spectrum decoding

An alternative to decoding on the raw signal at each point in time is to use both the real and imaginary parts of the complex-valued Fourier coefficients as features/inputs to a classifier. We will refer to this decoding paradigm as *complex spectrum decoding*. When all this information is included as features for classification, then the resulting information content in each frequency band is given by:

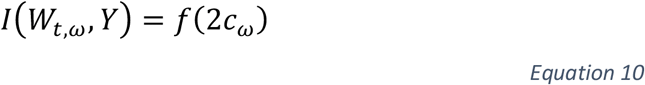

Where *c*_*ω*_ is the average value of the sinusoidal expression associated with the real information content in Equation 4 (see Appendix E for proof). Importantly, this expression is no longer sinusoidal; it is stable over time, and greater or equal to the peak information content that can be obtained using only the real spectrum (see Figure 4). Consequently, this overcomes both the problematic interpretational issues associated with *instantaneous signal decoding* discussed above, as well as the risk of representational aliasing that would otherwise require low-pass filtering with cut-off one quarter of the sampling rate.

**Figure 4:**
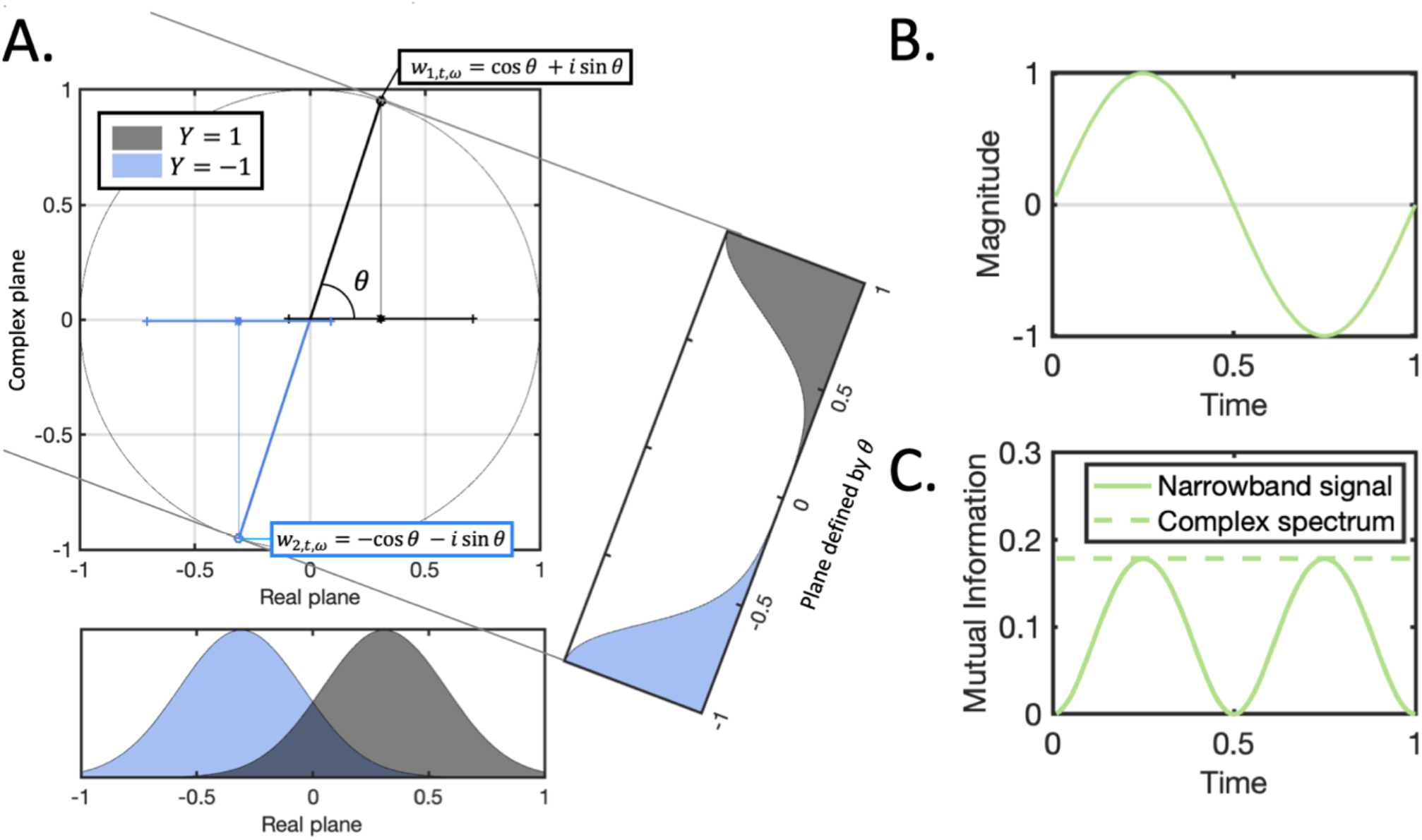
Motivation for complex spectrum decoding. **A**. The signal modelled by *w*_*n,t,ω*_ can be visualised as a point rotating around a circle in the complex plane, with the two stimulus conditions shown in grey and blue corresponding to opposite sides of the circle. As the signal crosses the imaginary axis (i.e. as *θ* approaches 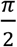), the separability of the two conditions in the real plane is minimised (corresponding to the troughs in the narrowband information content in Figure 2); however, at the same point, the two conditions in the complex plane (see the plane defined by *θ*) are still highly separable. In fact, projecting onto the plane defined by the instantaneous phase *θ* results in information content that is stable over time and not varying with the phase of the oscillatory signal. B. The real part of the signal (i.e. *z*_*n,t,ω*_ = *Re*(*w*_*n,t,ω*_)) is a sinusoid as previously characterised. C. The corresponding narrowband mutual information (i.e. *I*(*Z*_*t,ω*_, *Y*)) drops to zero in the sinusoidal troughs, whereas the complex spectrum information term (i.e. *I*(*W*_*t,ω*_, *Y*)) is constant throughout.

#### 3.2. Practical considerations for non-stationary and non-oscillatory signals

We emphasise the generality of these results, deriving from the fact that *any* arbitrary time series can be mapped into the frequency domain by a Fourier decomposition. Whilst we have so far simulated quite simplified evoked responses comprising only a few frequency components, our approach generalises to those that contain non-stationary and/or non-oscillatory components. In this section we demonstrate this with some more complex simulations.

##### 3.2.1. Sliding window Fourier decompositions

Real evoked responses are more complex than the illustrative examples we have simulated so far and in particular do not have spectral profiles that are constant over the whole trial epoch. We therefore anticipate that the methods introduced above will be most informative when combined with sliding window methods, e.g. where separate Fourier decompositions are applied to each window within a trial epoch rather than a single Fourier decomposition applied to the whole epoch.

There are numerous methods for estimating spectral properties over sliding windows, which are typically similar in motivation but different in implementation. Perhaps the most important factor is how the trade-off between time and frequency resolution is handled. Given our focus on characterising representational dynamics over time, we prefer methods that use a fixed temporal resolution, such as the Short-Time Fourier Transform (STFT). This provides complex-valued Fourier coefficients in each frequency band at each timepoint within a trial, allowing decoding accuracy to then be computed timepoint-by-timepoint without the interpretational problems previously discussed.

##### 3.2.2. Non-stationary oscillatory signals

To test these methods on evoked signals characterised by transient spectral properties, we simulated a signal over two channels using a combination of frequency chirp functions and unit step functions (example 1 in Figure 5). To maintain simplicity only one of the two conditions has this profile, the other is a null condition of stationary Gaussian noise. As shown by the time-frequency diagram on Figure 5A, the frequency distribution of the signal varies over time and over the two channels. For this signal, we then computed:

**Figure 5:**
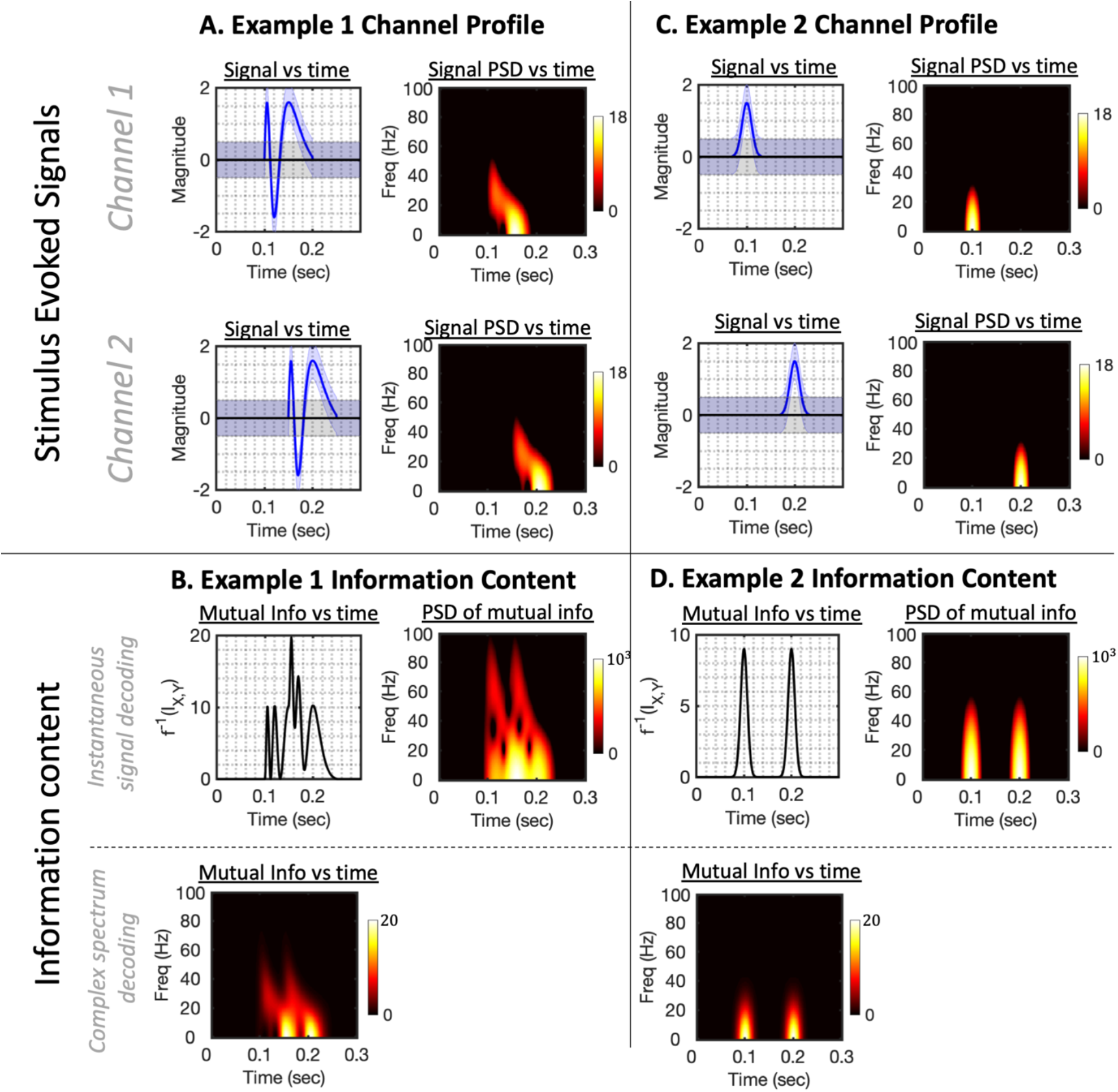
Complex spectrum decoding remains applicable and informative even when the spectral properties vary over time or are not fundamentally oscillatory. **A**. In Example 1, we simulate a signal across two channels with a time-varying frequency ‘chirp’ response, with a different onset time on each of the two channels. Left hand side plots the actual trial-averaged evoked response for each channel, right hand side the PSD as a function of time on each channel, showing the frequency content is transient on each channel and limited to frequencies below 50Hz. **B**. The information content associated with this signal. Top row plots the information content obtained by doing *instantaneous signal decoding*; right hand side plots the frequency profile of this mutual information timecourse, which reflects a mix of the spectrum from the two channels translated to double their original frequency (i.e. up to 80 Hz). Lower plot shows the mutual information obtained by *complex spectrum decoding* in each frequency band, which reflects the true frequencies at which information is present in the original signal. **C**. In Example 2, we simulate a non-oscillatory evoked response comprising two distinct processes occurring at different times; these characterise the signal over each channel in time and frequency. **D**. The information content available to instantaneous signal decoding identifies and separates these processing stages. This profile of two distinct peaks is similarly recovered from the complex spectrum information content (provided a suitable size of sliding window is used), demonstrating that this approach does not obscure such features where they are genuinely reflected in non-sinusoidal activity.

i. The *broadband information content;* This corresponds to the information content available to ***instantaneous signal decoding***, i.e. the timepoint-by-timepoint decoding approaches that are most typically used in the literature (T. Carlson et al., 2013; Cichy & Pantazis, 2017; Grootswagers et al., 2017).
ii. The *complex spectrum information content*; this corresponds to the information content available to ***complex spectrum decoding*** as we have proposed. In this case however we have estimated the complex spectral features using a sliding window (specifically using a STFT with 50ms sliding Hamming window).

As shown in Figure 5B, the broadband information content (analogous to the decoding accuracy obtained by *instantaneous signal decoding*) contains fast dynamics that do not clearly relate to the evoked signal shown in Figure 5A. Applying a similar STFT analysis to this information content (Figure 5B, right hand side) shows it reflects components at up to double the frequency of the corresponding signals (i.e. it contains components at up to 100Hz, double the frequencies identified in Figure 5A).

In contrast, the complex spectrum information content provides frequency band specific measures of information content that more closely reflect the spectral distribution of information at each moment in time over the course of the trial (i.e. Figure 5B, lower plot reflects the combined contributions of the channel power spectral density plots in Figure 5A). From the perspective of representational dynamics, such information is at least complementary, and we would argue more informative than that available to *instantaneous signal decoding*.

##### 3.2.3. Non-oscillatory evoked signals

In Results Section 1 we showed that consecutive peaks in decoding accuracy timecourses could arise due to a simple oscillatory signal, even if this oscillatory signal is itself stable over time. We argued that these peaks should not be interpreted as representing discrete events or cognitive phenomena. This begs the question, how do our methods perform if the underlying signals *do* derive from discrete temporal events, where the underlying signals cannot be parsimoniously represented using sinusoidal components?

To test this, we simulated an evoked response deriving from two spatially and temporally distinct “activations”, and repeated the analysis described above to compare the broadband and narrowband information content. To simulate non-oscillatory signals, each activation was characterised by a Gaussian kernel ssfunction (Figure 5C). As shown in Figure 5D, the broadband information content (i.e. that available when doing *instantaneous signal decoding*) produces two distinct peaks corresponding to each activation. Notably, this profile is replicated in the complex spectrum information content (Figure 5D, lower panel) showing that this method does not obscure such phenomena – provided the sliding window width is less than the period between these activations. Wider window lengths progressively include more information from both activations and the peaks become much less pronounced (see Supplementary Information Section S2 and Figure S2). We therefore conclude that, subject to appropriate sliding window sizes, *complex spectrum decoding* can eliminate the fast dynamics associated with sinusoidal components of the evoked response, whilst not eliminating the structure associated with spatially distinct, potentially non-oscillatory evoked activations.

### 4. Evidence from MEG data

The results we have presented are fundamentally theoretical and supported by simulated data from models of evoked activity. We therefore wanted to test how these findings extend to real data, and therefore tested our main predictions on a MEG dataset of visual image decoding.

We took a publicly available dataset comprising 15 subjects viewing 118 different visual stimuli (Cichy et al., 2016). The epoched data was then decoded (see Methods) to predict the trial condition labels using the three paradigms:

i. ***Instantaneous signal decoding***: decoding the raw broadband signal time-point-by-timepoint as widely performed in the literature (T. Carlson et al., 2013; Cichy & Pantazis, 2017; Grootswagers et al., 2017).
ii. ***Narrowband Signal decoding***: sliding window decoding using the time-frequency estimates from the STFT, but only using the real coefficients across all sensors as a set of features. This method is analogous to decoding on data filtered into specific frequency bands of interest.
iii. ***Complex spectrum decoding***: sliding window decoding using the time-frequency estimates from the STFT, using both the real and imaginary coefficients across all sensors as a set of features.

#### 4.1. Decoding accuracy vs time in different decoding paradigms

Figure 6A plots the decode accuracy derived from decoding under the three identified paradigms. As paradigms (ii) and (iii) provide accuracy in each frequency band independently, for ease of visualisation they are each plotted separately against paradigm (i). Averaged over all pairs of stimuli and all subjects, this identifies a systematic variation in the information content at different frequencies as a function of time. The earliest detectable information appears in higher frequencies, but these peak quite transiently at relatively low values and are quickly surpassed by information content in lower frequencies, which rise to higher values and are then sustained for a longer duration. Notably, the information in either the 10Hz or the 0Hz band exceeds that obtained by *instantaneous signal decoding* for nearly the entire period analysed. From the perspective of representational dynamics, this establishes first and foremost that Fourier decompositions can improve decoding accuracy over *instantaneous signal decoding* methods whilst retaining a profile of how the representational content evolves in both time and frequency.

**Figure 6:**
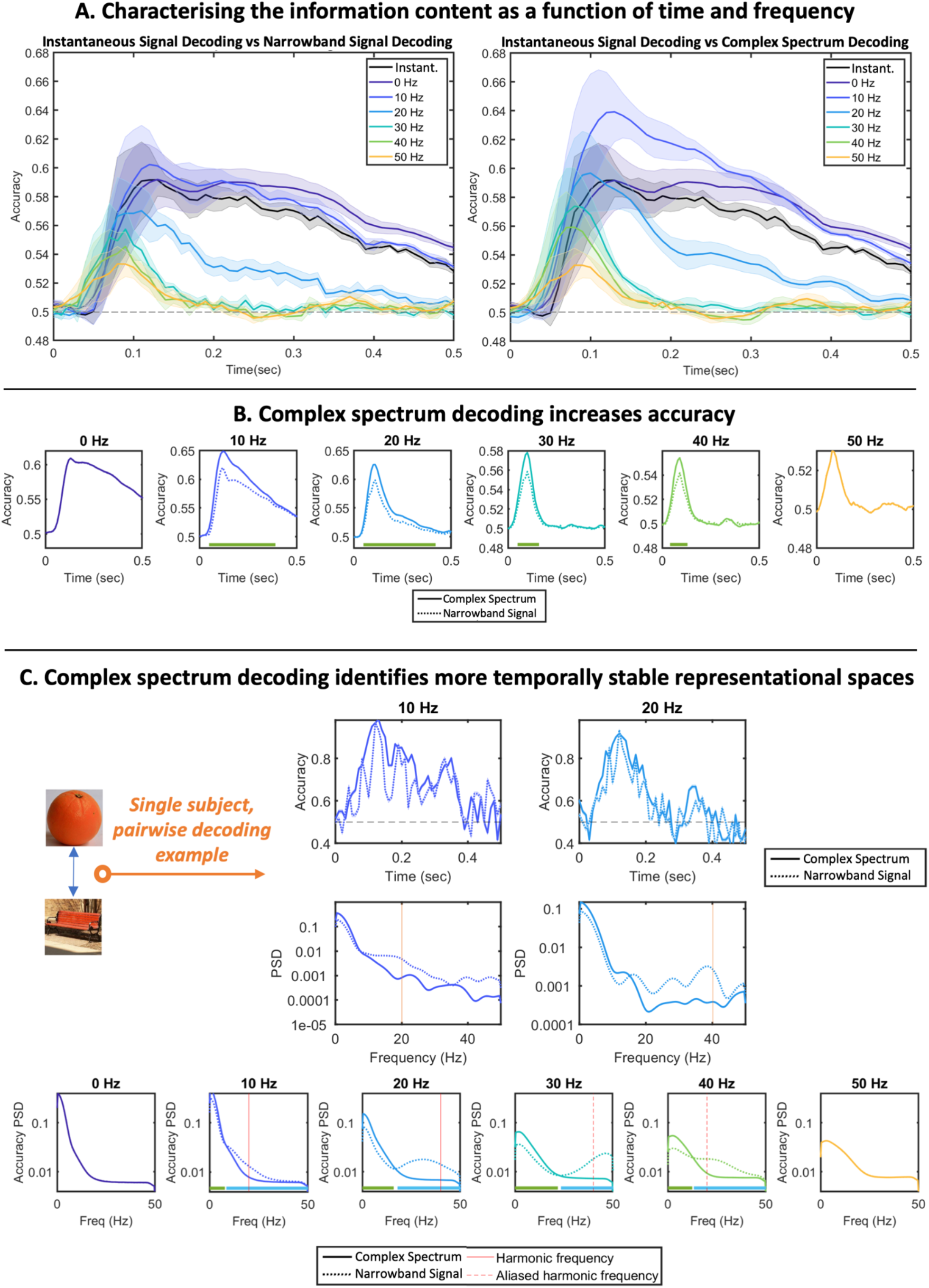
Characterising information content across the frequency spectrum. **A**. Decoding accuracy timecourses obtained by *instantaneous signal decoding* vs either *narrowband signal decoding* (left) or *complex spectrum decoding* (right). All plots show mean +/-SE over subjects. **B**. Directly comparing the accuracy vs time in each frequency band of *narrowband signal decoding* and *complex spectrum decoding*. Significance bars denote periods where *complex spectrum decoding* accuracy is significantly greater than *narrowband signal decoding*: p<0.01 using cluster permutation tests. **C**. Single subject results. The dynamics, and therefore any attempt to assess the temporal stability of the decode accuracy, are obscured by averaging over all participants and subjects. Top: taking a single subject, single pairwise comparison as an example, we show the decode accuracy obtained by either *narrowband signal decoding* or *complex spectrum decoding* as a function of time (upper) or frequency (lower). These show significant harmonic components at double the fundamental frequency in each band when *narrowband signal decoding* is used. Lower plot: We took all individual accuracy vs time plots (across all subjects and all stimulus comparisons) and computed their power spectral density, then averaged (plots show mean +/-SE over subjects). The power spectrum obtained using *narrowband signal decoding* show strong peaks at harmonic frequencies (for 10Hz and 20Hz bands) and at aliased frequencies (for 30Hz and 40Hz bands). Significance bars denote significance at p<0.01 levels using cluster permutation tests; green bars denote *complex spectrum decoding* greater than *narrowband signal decoding*; blue bars denote *narrowband signal decoding* greater than *complex spectrum decoding*. This shows that, in all frequency bands, the increased accuracy obtained by *complex spectrum decoding* is concentrated in lower frequencies, reflecting more temporally stable representational spaces.

#### 4.2. Complex spectrum decoding accuracy exceeds narrowband signal decoding accuracy

Figure 6B compares the average classification accuracy in each frequency band, averaged over all subjects and pairs of stimuli, when either the *complex spectrum decoding* or *narrowband signal decoding* is applied (it follows from the definition of the discrete Fourier transform that the imaginary coefficients in the 0Hz and 50Hz frequency bands are always zero, so in these bands the two paradigms are in fact equivalent). In all cases the classification accuracy obtained using *complex spectrum decoding* exceeds that obtained using *narrowband spectrum decoding*; this information gap can be interpreted as the information stored in the gradient of these sinusoidal components.

#### 4.3. Narrowband signal decoding produces spectral peaks at double their original frequency in inferred decoding accuracy metrics

Our models predict that the information content associated with evoked spectral components at a given frequency is itself oscillatory at double that frequency, unless *complex spectrum decoding* is applied. We have so far plotted the average over all subjects and all comparisons, therefore obscuring some of the temporal dynamics evident in each comparison. For example, in Figure 6C we plot the timecourse obtained for one subject and one pair of stimuli; the accuracy timecourse obtained from *complex spectrum decoding* appears to follow the envelope of the equivalent timecourse obtained by *narrowband signal decoding* which appears to show sinusoidal dynamics. If we take the PSD of these accuracy timecourses, we observe a peak at double the frequency band being analysed (i.e. the 10Hz and 20Hz bands are associated with a 20Hz and 40Hz spectral peak respectively). If we take the PSD of the timecourse for every pair of stimuli and every subject and average, we see the PSD is significantly higher in the 10Hz and 20Hz bands at approximately 20Hz and 40Hz, respectively.

Given a sampling rate of 100Hz, we expect representational aliasing to occur linked to any evoked spectral content above 25Hz. Specifically, given evoked spectral content at 30Hz or 40Hz, we expect representational aliasing artefacts at 40Hz and 20Hz, respectively (for example, a 30Hz component is translated to 60Hz in the accuracy timecourses; as this is 10Hz above the Nyquist frequency, it is aliased to 10Hz below the Nyquist frequency, i.e. to 40Hz). For both of these narrowband signals, we see peaks at these locations, confirming the presence of representational aliasing. We stress that this aliasing effect must also be present in the *instantaneous signal decoding* results, they just cannot be explicitly resolved as we have no knowledge of the frequencies at which they would be expected.

Finally, in these plots we note that spectra are significantly more weighted towards the lower end of the frequency spectrum for *complex spectrum decoding* vs *narrowband signal decoding*, whilst the opposite relationship is the case towards the upper end of the frequency spectrum. This means that the higher accuracies obtained by *complex spectrum decoding* in Figure 5B are a result of increased low frequency content, or representational dynamics that are more stable over time.

#### 4.4. Complex Spectrum Decoding accesses information content that is complementary over frequencies

Having established that *complex spectrum decoding* accesses information content that is not available to *instantaneous signal decoding*, one final question arises; is the complex spectral information across different frequencies overlapping, or complementary? That is to say, if we aggregate the information over frequency bands, do we obtain performance that is merely equivalent to the best individual frequency band – or exceeding it? To answer this question, we trained an aggregate classifier to estimate the aggregate information distributed over all frequency bands (see Methods).

Figure 7A plots the performance of the aggregate classifier against the *complex spectrum decoding* accuracies achieved in each frequency band, and that obtained by *instantaneous signal decoding*. The aggregate classifier significantly outperforms the *instaneous signal decoder*, reaching a peak accuracy of 67.6% vs 61.6%. As plotted in Figure 7B, this difference quantifies the total amount of information that is inadvertently being omitted by the insensitivity of *instantaneous signal decoding* paradigms to information stored in signal gradients. However the aggregate decoder accuracy also peaks at a level higher than that obtained in *any* individual frequency band. As in Figure 7C, over the period between 70msec and 190msec following stimulus presentation, the aggregate classification accuracy significantly exceeded the information content in any individual frequency. This coincides with the time over which significant information content was distributed across multiple frequency bands, especially higher frequency bands, proving that these different frequency bands contain information content that is complementary. The performance is quite different for timesteps more than 370msec after stimulus onset, with the ensemble classifier underperforming slightly relative to the best narrowband classifiers (albeit still outperforming standard broadband methods). Over this period, the classifiers trained on higher frequencies output chance level predictions, and only lower frequency bands contain meaningful information content. We conclude that over this period, all meaningful information is concentrated in lower frequency bands, and the inclusion of high frequency bands that only contain noise is in fact detrimental to classifier performance.

**Figure 7:**
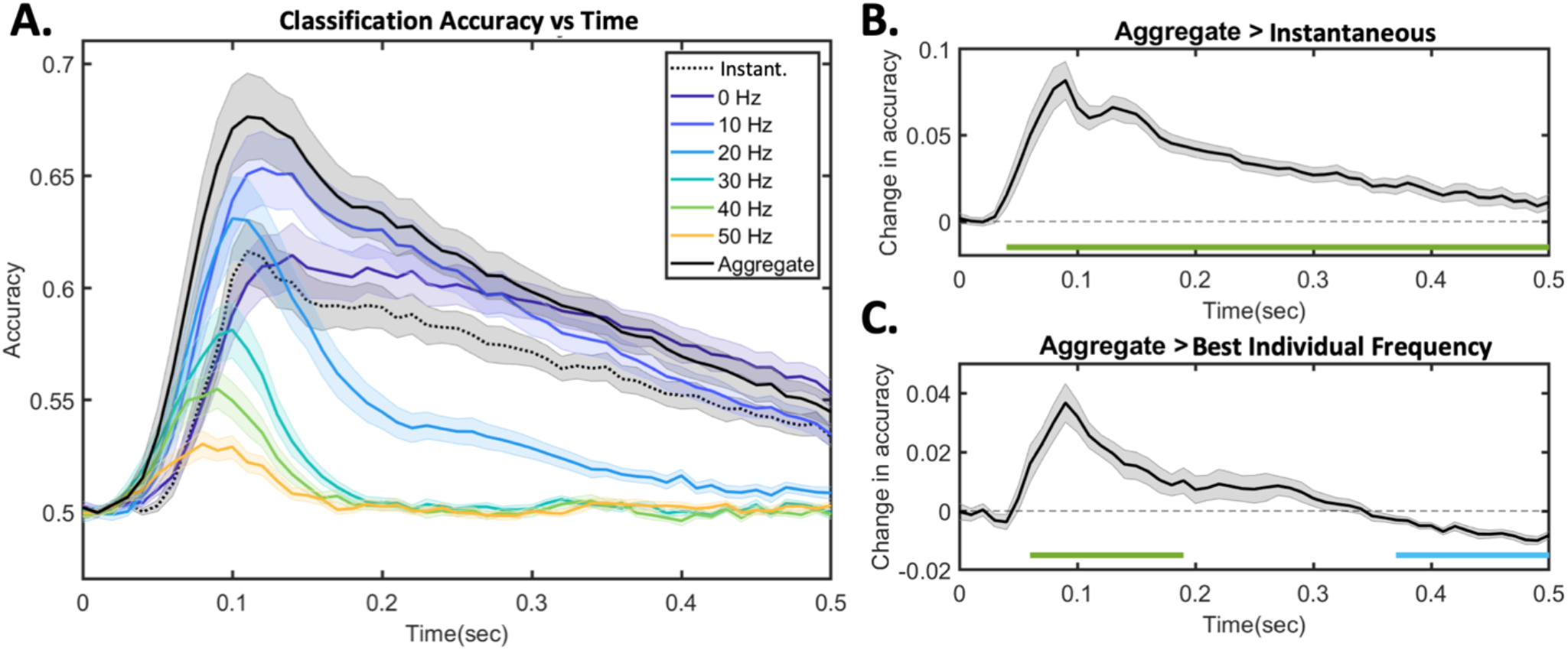
Classification accuracy obtained by aggregating information over multiple frequency bands. **A**. Plotting the classification accuracy achieved vs time for *instaneous signal decoding*, for each individual frequency using *complex spectrum decoding*, and then for the aggregate classifier. All plots show mean +/-SE over subjects. **B**. Plotting the contrast of the aggregate classifier accuracy minus the instantaneous signal decoding accuracy; significance bars denote p<0.01 using cluster permutation tests, green bars denote contrast is significantly positive, blue bars denote the sontrast is significantly negative. **C**. Plotting the contrast of aggregate classifier accuracy minus the best individual frequency (i.e. the highest accuracy obtained by any *complex spectrum decoder* at each timepoint); significance bars defined as for B.

## DISCUSSION

We have outlined a widely overlooked problem in decoding pipelines: that frequency components in the evoked response produce corresponding components at double their original frequency content in the resulting accuracy metrics. Where researchers are not aware of this fundamental relationship, there is a considerable risk of misinterpreting results and, in particular, of inferring relationships with canonical frequency bands that are in fact trivial representations of the evoked response spectrum.

Our results are fundamentally mathematical, and should be interpreted as such; they derive from the expected Fourier spectrum of the evoked response, not from the fundamental frequency of a canonical neural oscillation. For example, a 40Hz Fourier component can be produced by a vast range of underlying neural sources, only a subset of which would be considered ‘gamma oscillations’. Our results apply regardless; thus the recommendation to low-pass filter with a cut-off frequency of one quarter of the sampling rate applies to any researcher doing *instantaneous signal decoding*, irrespective of the frequencies of neural activity they may be interested in or expecting.

As an information theoretic result, if our modelling assumptions hold then these results are fundamental and apply to any *instantaneous signal decoding* approach regardless of methodological choices on the part of the researcher; they cannot be overcome by use of nonlinear classifiers, machine learning tools, or by analysing different accuracy metrics. In our analysis we have derived the spectrum of the information content up to an arbitrary monotonic scaling denoted by the function *f*. It follows that other widely used metrics to assess decoding accuracy (such as classification accuracy, distance from the classification hyperplane etc.) are each a different monotonic scaling of this quantity (see table 2 and SI for further details). We therefore argue that our results are universally applicable to *instantaneous signal decoding* pipelines regardless of any variations in methodological choices.

**Table 1:** Overview of random variables modelled in this paper.

| Random Variable | Observation on nth trial | Domain and dimension | Used to model: |
| --- | --- | --- | --- |
| $X_t$ | $x_{n,t}$ | $\mathbb{R}^{1 \times P}$ | Recorded broadband signal at time $t$; input used for *instantaneous signal decoding* |
| $Y$ | $y_n$ | $\{1, -1\}$ | Stimulus class |
| $Z_{t,\omega}$ | $z_{n,t,\omega}$ | $\mathbb{R}^{1 \times P}$ | Narrowband signal at time $t$ in frequency band $\omega$; input used for *narrowband signal decoding* |
| $W_{t,\omega}$ | $w_{n,t,\omega}$ | $\mathbb{C}^{1 \times P}$ | Complex spectrum signal at time $t$ in frequency band $\omega$; input used for *complex spectrum decoding* |

**Table 2:**
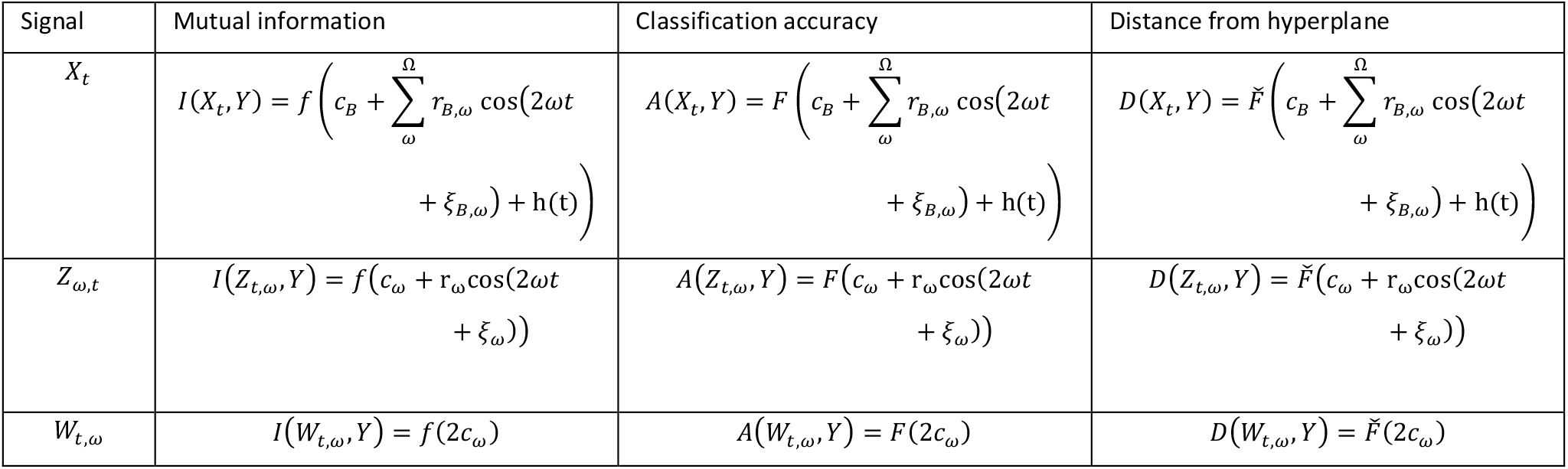
Mutual information results generalise to other common decoding metrics. We have characterised the information content associated with three variables up to an arbitrary monotonic function *f*. All our results generalise to other commonly used decoding metrics such as classification accuracy and distance from the hyperplane simply by substituting *f* by another monotonic function, here denoted by *F* and 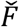 (see Supplementary Information Section S1 and Figure S1 for plots of these functions).

We have characterised three major decoding paradigms but do not claim these to be exhaustive with respect to the literature. A very common approach involves the application of classifiers not to a recorded signal itself but to a set of Fourier features derived from a signal, which in most applications will be equivalent to the narrowband or complex spectrum decoding paradigms such that all our results remain applicable. However an emerging area of research infers nonlinear time-domain features, for example through the training of temporal convolutional networks or recurrent neural networks, that are then used as inputs for classification (Kalafatovich et al., 2020; Schirrmeister et al., 2017; Zubarev et al., 2019). These methods typically offer a greater ability to separate conditions, however the accompanying barriers to interpretability have to date limited their direct application in the study of representational dynamics. We hope that such interpretability barriers will be challenged and overcome in future work, and that the relationships we have outlined here may aid this endeavor.

Finally, we have shown that *complex spectrum decoding* overcomes the problem of representational aliasing whilst also presenting other benefits; specifically, leading to higher accuracies that are more stable over time. However, it similarly presents its own challenges. The significant increase in dimensionality associated with a feature vector that varies simultaneously over time, space and frequency may present computational challenges. Furthermore, whilst we see interpretational benefits to having results that are resolved in both frequency and time, in some circumstances (such as the non-sinusoidal signal example simulated in Results Section 3.2.3) this additional complexity may not harbour any new insights. We have spoken broadly of Fourier analysis, again to stress that these results apply generically to STFTs, wavelet decompositions, or any other such method – however each of these apply different assumptions that mostly result in different trade-offs of time and frequency resolution. These trade-offs are likely to be especially pertinent in the context of high temporal resolution decoding. Nonetheless, the benefits can be quite substantial and well justified by the results.

In conclusion, we have characterised the relationship between the stimulus evoked spectrum and the information content spectrum, which is commonly used to investigate the brain’s representational dynamics. Understanding how these two quantities relate is crucial to interpreting results obtained via decoding pipelines. By establishing these relationships under three different decoding paradigms, this work opens the door to much stronger interpretation of decoding results by linking the question of *what* is being represented with the neural mechanisms explaining *how* it is being represented. We hope this will enable more targeted scientific enquiry to uncover the true mechanisms by which the brain processes diverse forms of information.

## METHODS

### 1. Model outline

The assumptions expressed in Results Section 1.1 can be more fully expressed mathematically, with corresponding expressions obtained for the probability distribution of each of the random variables *X*_*t*_, *Y* and *Z*_*t,ω*_.

We have assumed that the stimulus class is binary with equal class probabilities. This corresponds to the following distribution for *Y*:

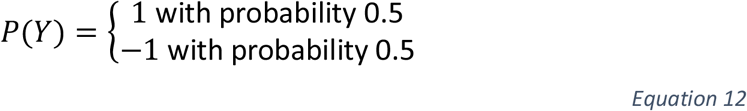

Following on from Equation 1, since the *μ*_*t*_ term captures the mean over both conditions, we can then model the expected value of each signal *z*_*n,t,ω*_ on individual trials with polarity determined by the trial condition *y*_*n*_. We assume these narrowband signals have multivariate Gaussian noise that is independent and identically distributed (over trials and conditions) – this corresponds to an assumption that only the evoked response, and not the induced response, differs over the two conditions. This is a simplifying assumption that we discuss further and ultimately relax in Results Section 1.4. This allows us to specify the probability distributions of *Z*_*t,ω*_ (of which *z*_*n,t,ω*_ is the sample corresponding to the *n*th trial) as follows:

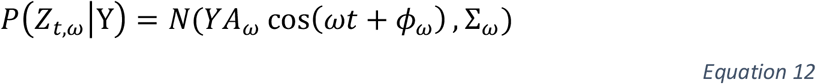

Each *A*_*ω*_ term is a diagonal [*P* × *P*] matrix, where the *i*th diagonal entry, denoted by *a*_*ω*,V_, reflects the magnitude of the component at frequency *ω* on channel *i*. Both *ω* and *t* are scalar indices reflecting the frequency and time respectively, whereas *ϕ*_*ω*_ is a [*P* × 1] vector, each entry of which contains the phase offset of the oscillation at frequency *ω* across the *P* channels. Finally, we model induced effects (i.e. narrowband power that is not phase aligned to the stimulus) independently in each frequency band, where Σ_*ω*_ is the [*P* × *P*] covariance matrix modelling the spatial variance and correlations expressed at frequency band *ω*.

For any set of discretely sampled data recordings with at least *P* total trials (i.e. more trials than channels), all of the above parameters are fully identifiable. The data for each channel and each trial can be decomposed into a discrete Fourier series representation of the above form where the number of frequency components equals half the number of timepoints in the trial 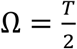 (for simplicity we here model a single Fourier decomposition over the trial; this can equivalently be computed over sliding windows as in Results Section 3, in which case the number of frequency components equals half the number of samples in a window). Following from the uniqueness of the Fourier transform, unique values can be obtained for the diagonal matrix *A*_*ω*_, the phase offsets *ϕ*_*ω*_ and the patterns of spatial correlations in each frequency Σ_*ω*_.

The distribution of the broadband signal *X*_*t*_ given in Equation 2 then follows directly from Equation 1 (by observing that a sum of narrowband Gaussian components is also Gaussian distributed).

### 2. MEG decoding methods

We took a publicly available dataset comprising 15 subjects viewing 118 different visual stimuli (Cichy et al., 2016). This data had been acquired on an Elekta Neuromag scanner with 306 channels (204 planar gradiometers and 102 magnetometers) at 1kHz sampling rate, with filtering applied at acquisition with bandpass 0.03Hz to 300Hz. We downsampled the data to 100 samples per second with an anti-aliasing filter with cut-off at 50Hz and extracted the 0.5 second epochs immediately following stimulus presentation. The data was then mapped into a complex time-frequency decomposition using an STFT with Hamming window length of 100ms. The epoched data was then decoded to predict the trial condition labels using the three paradigms outlined in the text. Each approach fit linear support vector machine classifiers using three-fold cross validation. This was applied to each pair of the 118 images in a mass pairwise classification paradigm as originally implemented by (Cichy et al., 2016). In cases (ii) and (iii), classifiers were trained separately on each frequency band. The decoding used three-fold cross-validation to obtain independent classification accuracy metrics as a function of time and frequency for each pair of images and each participant.

Finally, to test the hypothesis that different frequency bands contained complementary information, we trained an aggregate classifier to estimate the aggregate information distributed over all frequency bands. We did this through a nested cross validation procedure. An inner cross validation loop simply consisted of the *complex spectrum decoding* estimates described above. The outer cross validation loop then partitioned all of the stimuli into two equally sized groups and applied two-fold cross validation to obtain accuracy estimates. This outer loop consisted of a random forest ensemble classifier with 100 trees, trained to predict the class label from the outputs of the *complex spectrum decoding* classifiers on each trial. This outer loop was run ten times with replacement for each subject, randomly sampling a different subset of stimuli with replacement on each cross validation fold.

## Appendices

### A. Mutual information for a Gaussian mixture model with equal covariances

Let us first consider a simpler model and derive a general result that we can then use to prove our claims. Suppose we have a random variable *Y* distributed as given in the text:

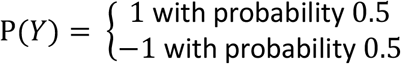

Suppose we then have another random variable *B* of dimension *P* × 1 conditioned on Y as follows:

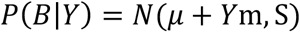

This is equivalent to a Gaussian mixture model with two components (corresponding to the cases *Y* = 1 and *Y* = −1) and equal covariances. The marginal distribution can then be expressed as follows:

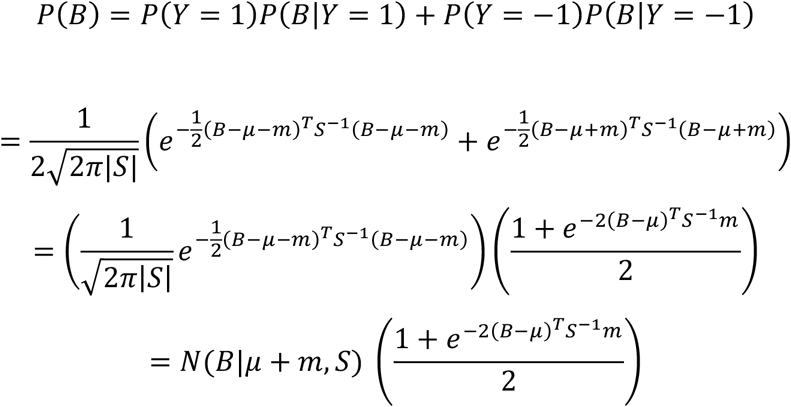

Where we use the notation *N*(*B*|*μ* + *m, S*) to denote the Gaussian distribution over *B* with mean *μ* + *m* and covariance *S*. We shall furthermore use the notation 𝔼_P(x)_*f*(*x*) to denote the expectation of a function *f*(*x*) given the probability distribution *P*(*X*). We can now compute the following result for the entropy of *B*:

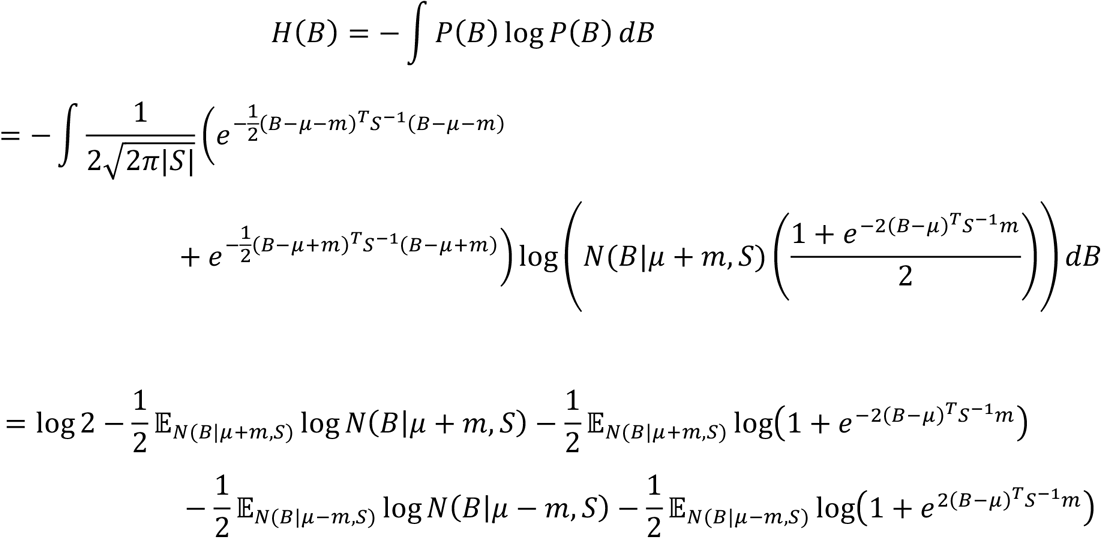

We then observe that the second and fourth terms correspond to the entropy of a multivariate Gaussian, which has a known solution; we similarly observe that the third and fifth remaining terms are each an expectation of a univariate function in *u* = 2(*B* − *μ*)^T^*S*^−1^ *m* and *v* = 2(*B* + *μ*)^T^*S*^−1^*m* respectively. With a substitution of variables this simplifies to:

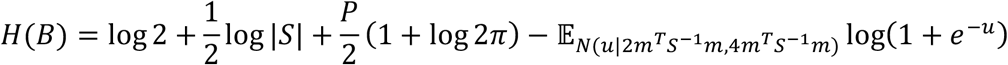

Similarly, we find that the conditional entropy is given by:

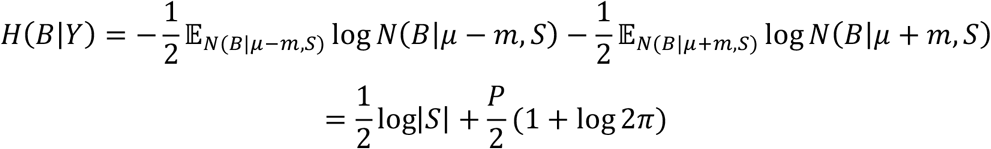

We can therefore apply the chain rule to derive the mutual information:

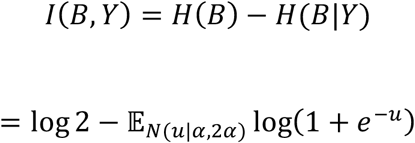

Note that the second term involves an integral that is intractable, but is a function of the scalar product *α* = 2*m*^T^*S*^−1^*m*. We therefore can state equivalently that:

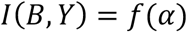

Where

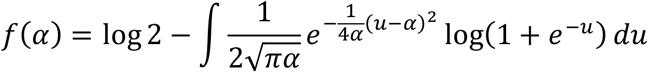

### B. Proof that the function *f* is monotonic and concave

Firstly, let us define the following probability distribution:

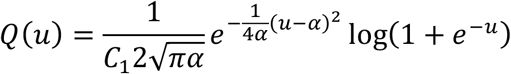

Where 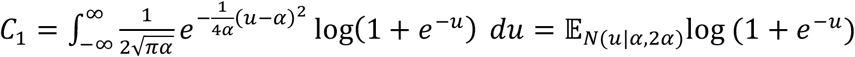. Our proof below rearranges the first and second derivative of *f* in terms of the higher moments of this distribution, thus we now seek an expression for these moments. The moment generating function for *Q*(*u*) is:

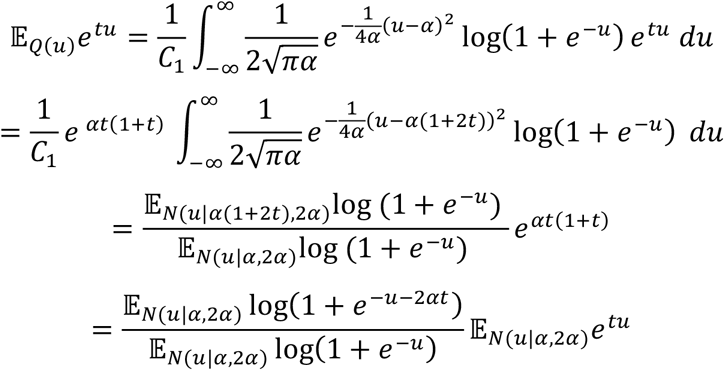

The moment generating function allows us to compute the higher moments of the distribution that are ultimately required for the proof. As the algebra for this is somewhat tedious we refer readers to Supplementary Information Section S3 for full details, where we obtain the following expressions for these moments:

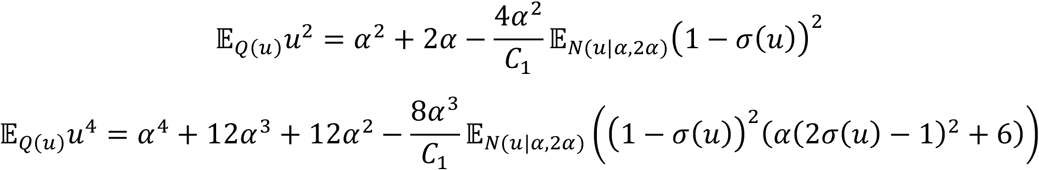

Where 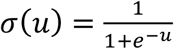 denotes the logistic sigmoid function. Note that the terms inside the expectations are strictly positive, such that their expectation is always greater than zero.

Now, consider the function *f*(*α*), plotted in Figure A1 and specified as in Appendix A.

**Figure A1:**
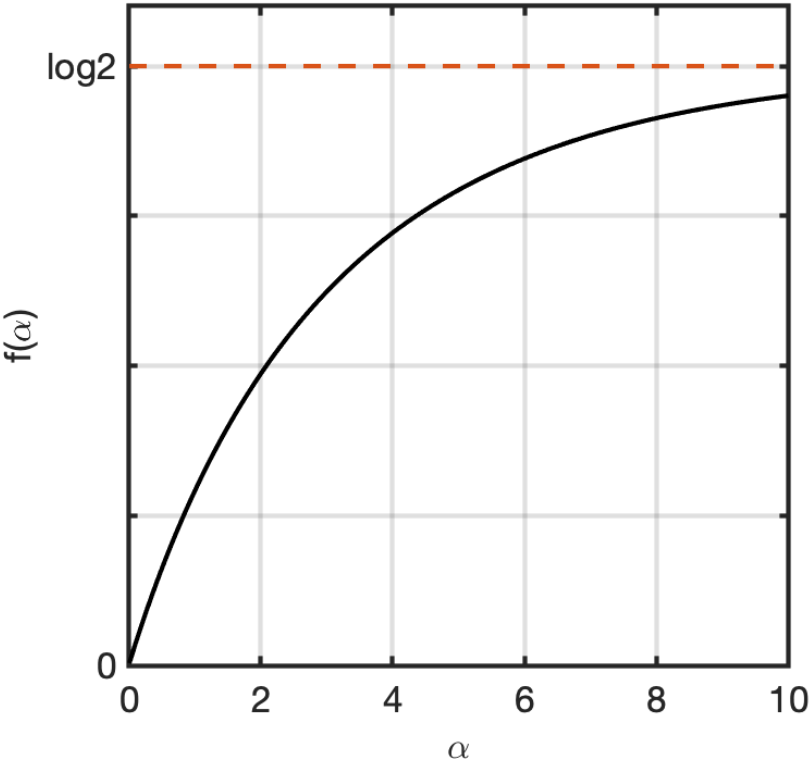
*the function f*(*α*), *which uniquely determines the information content of data generated from the Gaussian model specified in the text, is a monotonic concave function*.

The first derivative is:

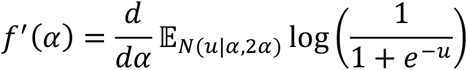

Let us denote by 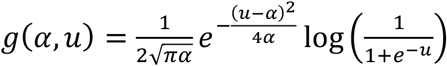. We can then evaluate by Leibniz rule:

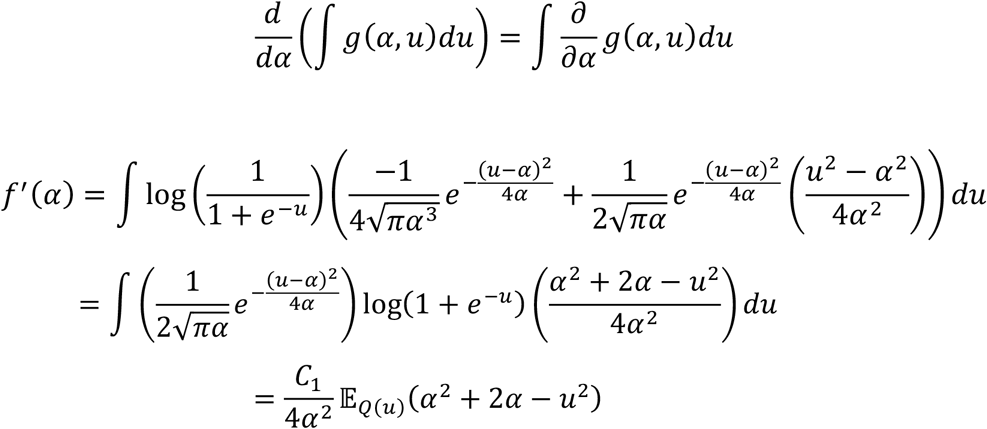

Substituting the above expression for 𝔼_Q(u)_*u*^2^, we have:

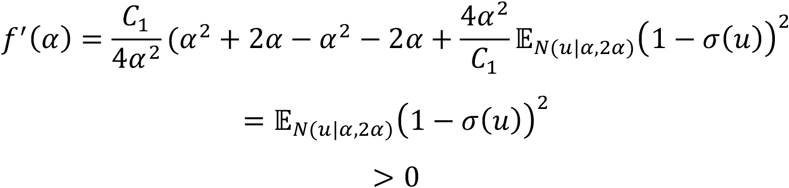

We conclude that *f* is monotonic.

A second application of Leibniz’ rule gives us the second derivative:

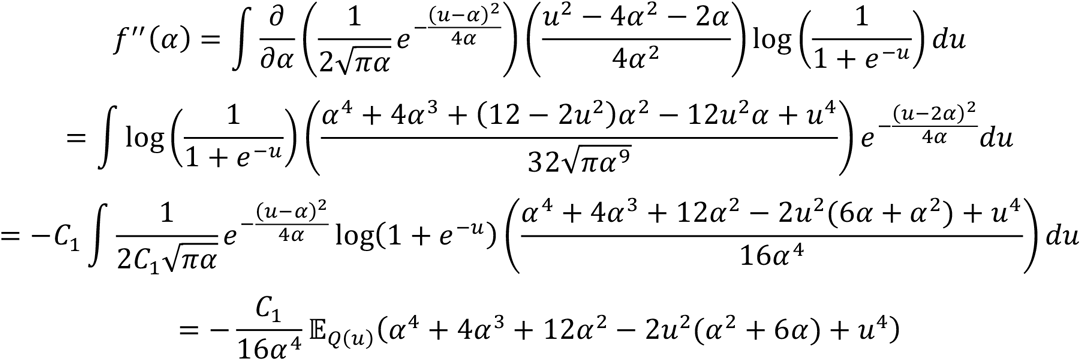

Now substituting the above expressions for 𝔼_Q(u)_*u*^2^ and 𝔼_Q(u)_*u*^4^ we again find that most terms cancel out leaving us with:

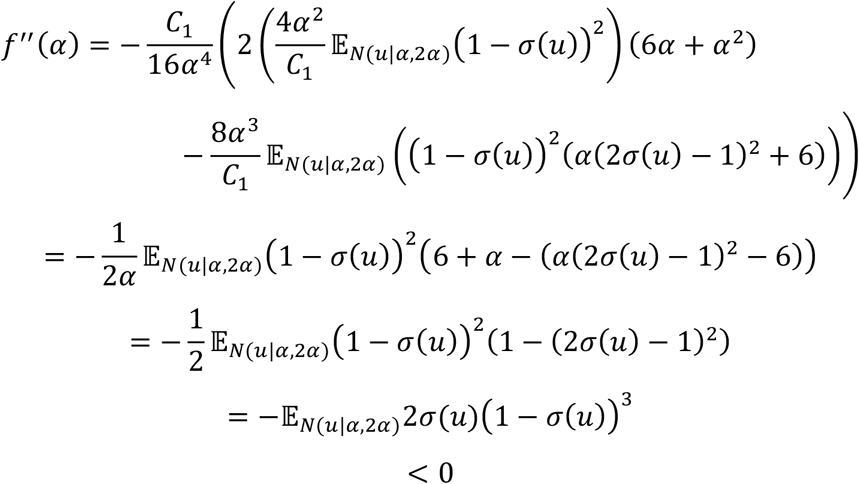

We conclude that *f* is concave.

### C. Mutual information of the narrowband signal

Consider the narrowband real signal *Z*_*ω,t*_ as specified in Equation 12. It can be seen that this is a special case of the model specified in Appendix A by substituting the following:

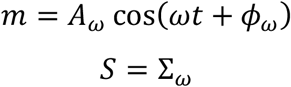

It therefore follows that the information content is given by:

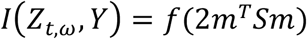

Re-writing the mean term in cartesian form, we have:

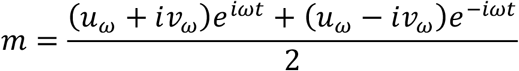

Where *u*_*ω*_ = *A*_*ω*_cos (*ϕ*_*ω*_) and *v*_*ω*_ = *A*_*ω*_sin (*ϕ*_*ω*_). This allows us to determine the information term:

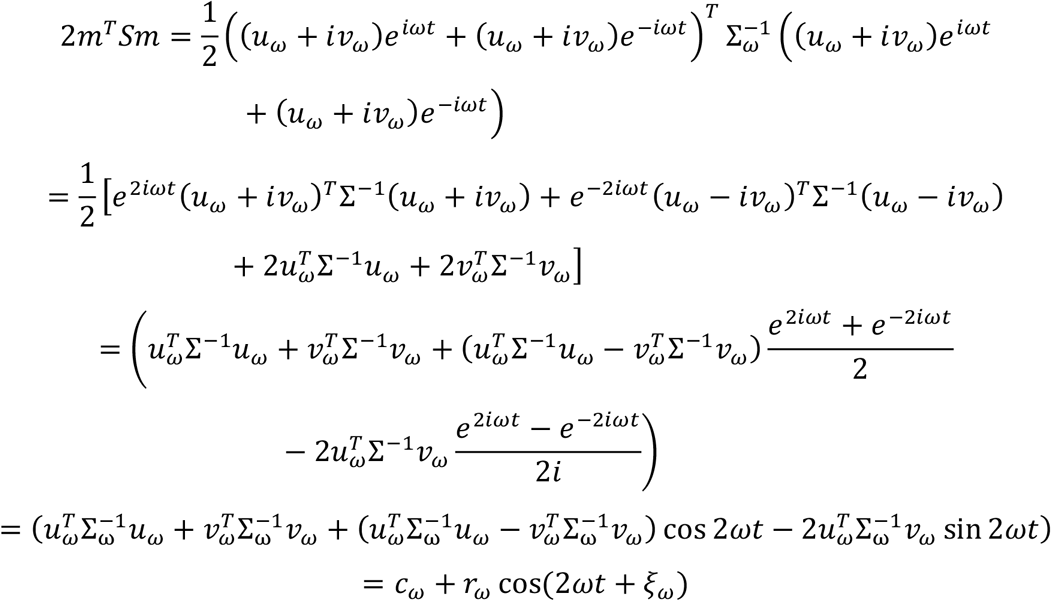

Alternatively, returning to the polar coordinates used throughout the paper, we have:

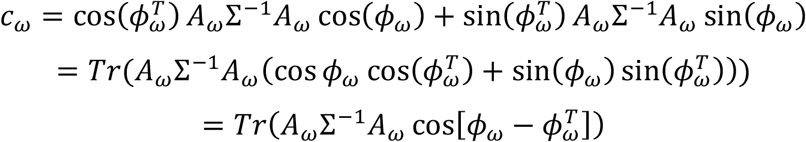

And applying the same steps for the remaining variables gives us:

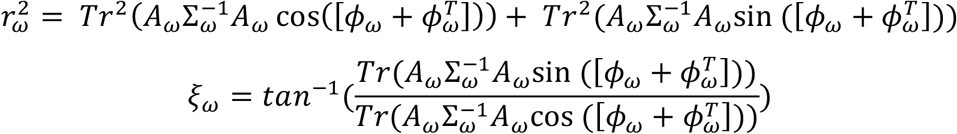

Where we have used the notation [*ϕ* ± *ϕ*^T^] for the [*P* × *P*] matrix constructed from the [*P* × 1] vector *ϕ*_*ω*_ such that the *i, j*th matrix entry is given by *ϕ*_*ω*,i_ ± *ϕ*_*ω*,j_. We have similarly used *vec* to denote the vec operator.

We conclude that the narrowband signal information content is given by:

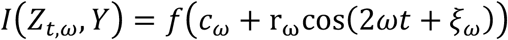

### D. Mutual information of the broadband signal

Consider the broadband signal *X*_*t*_ as specified in Equation 2. It can be seen that this is a special case of the model specified in Appendix A by substituting the following:

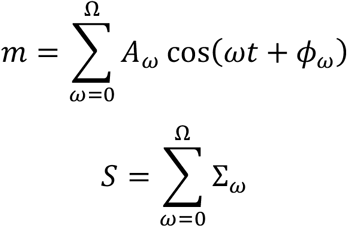

It therefore follows that the information content in the broadband signal is given by

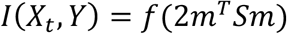

Substituting the above values:

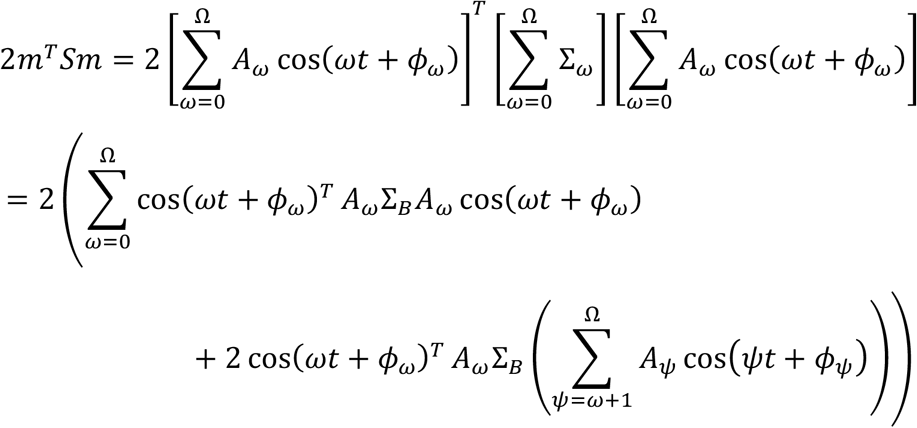

Writing in cartesian coordinates, such that:

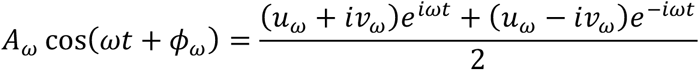

This becomes:

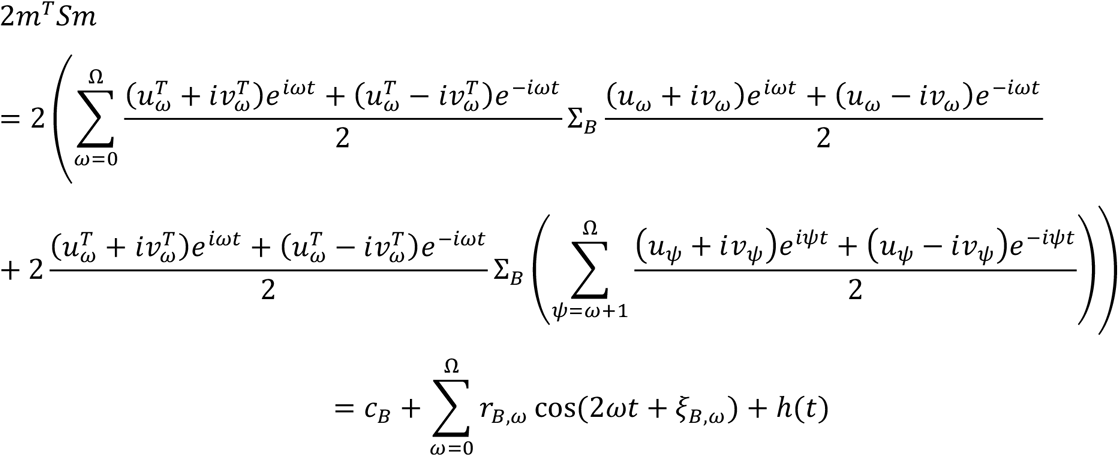

Where 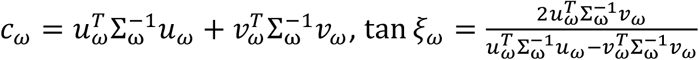 and 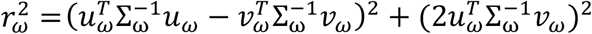, and the timeseries *h*(*t*) contains harmonic components at the sum and difference of each broadband frequency component:

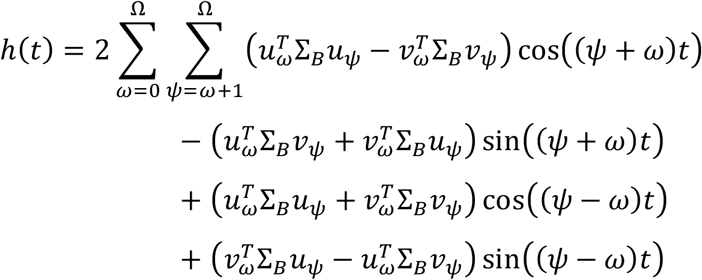

We conclude that the broadband information content is given by

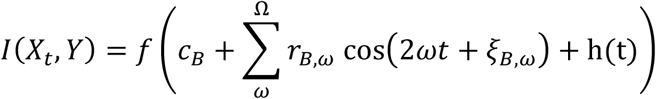

### E. Mutual information of the complex-valued Fourier signal

Consider the complex signal *W*_*ω,t*_ as specified in Equation 8. It can be seen that this is a special case of the model specified in Appendix A by substituting the following:

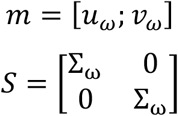

It therefore follows that the information content is given by:

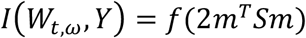

Substituting the above values we observe that:

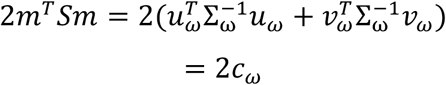

We conclude that the information content in the complex signal *W*_*ω,t*_ is given by:

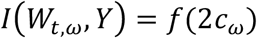

### F. Mutual information for a Gaussian mixture model with different covariances

The modelling above assumes that the induced response (i.e. the changes in power that have no phase alignment to the stimulus) has the same distribution over both stimulus conditions. This corresponds to an assumption that the narrowband covariance matrix is invariant over stimulus conditions. To explore how our results generalise to the case of stimulus-specific induced effects, let us return to the result of Appendix A and now define a new random variable 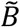 of dimension *P* × 1 as follows:

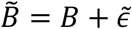

Where the new residual terms have distribution 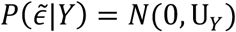 will be used to model induced effects. We assume that 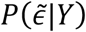 is independent of 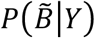. We previously defined *B* as a Gaussian mixture model with two components (corresponding to the cases *Y* = 1 and *Y* = −1) and equal covariances; thus the new random variable has a distribution given by:

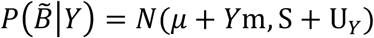

Which corresponds to a Gaussian mixture model with some common covariance given by *S* as well as some stimulus-specific covariance given by *U*_Y_. The mutual information for this variable is not tractable, however we instead obtain an upper bound by observing that 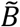 is a linear function of *B* and 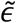, therefore the data processing inequality tells us that:

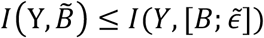

Where 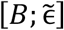 is the vector obtained by concatenating the random variables *B* and 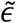. We shall now obtain an expression for the mutual information between this new term and the stimulus labels *Y*. Note that the marginal distribution is given by:

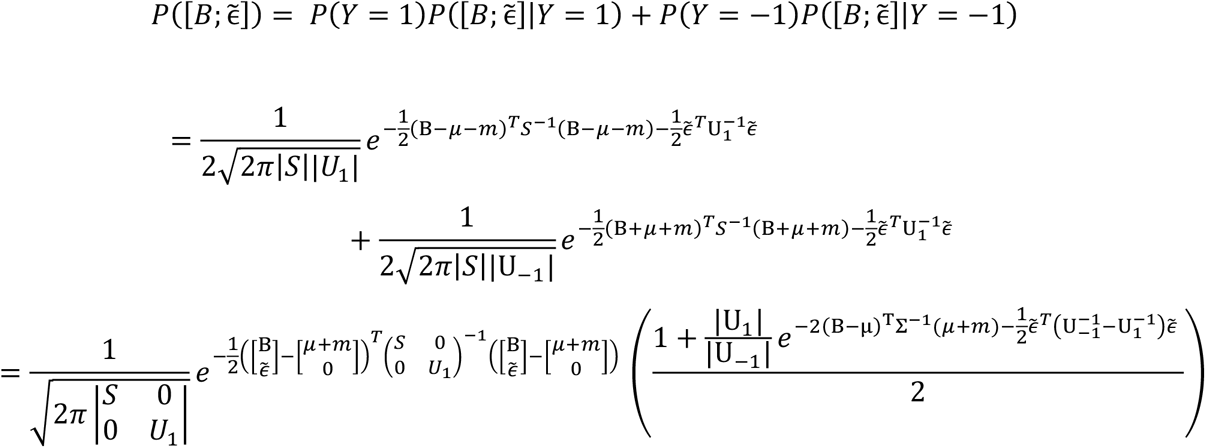

Following the same approach of Appendix A, we obtain the following for the entropy:

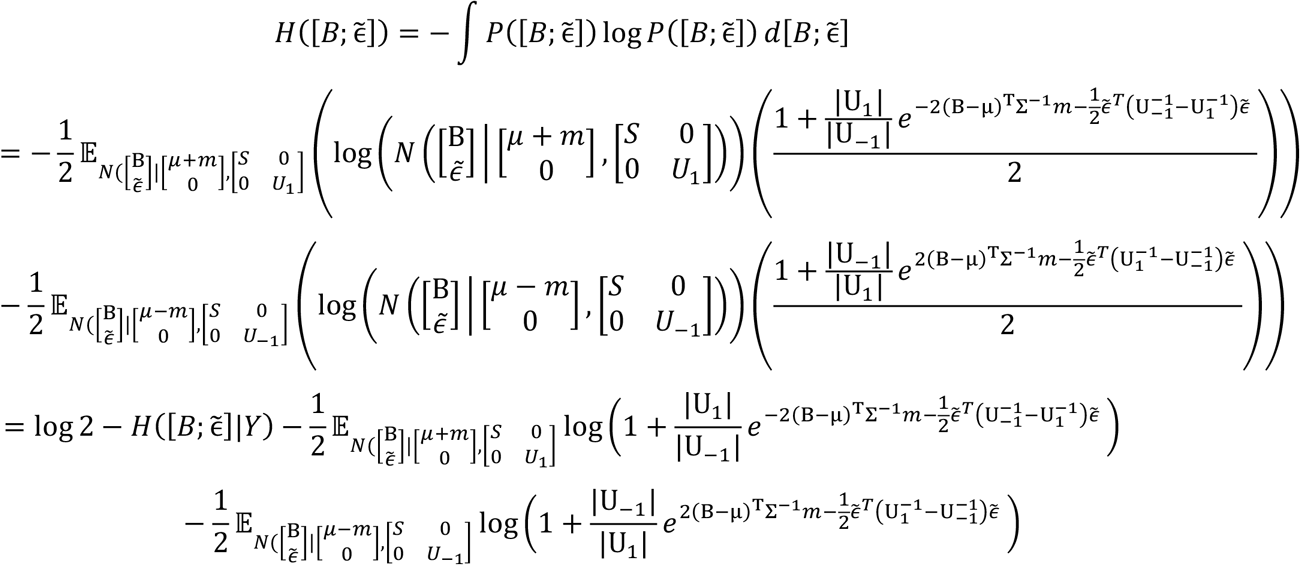

Applying the chain rule we have:

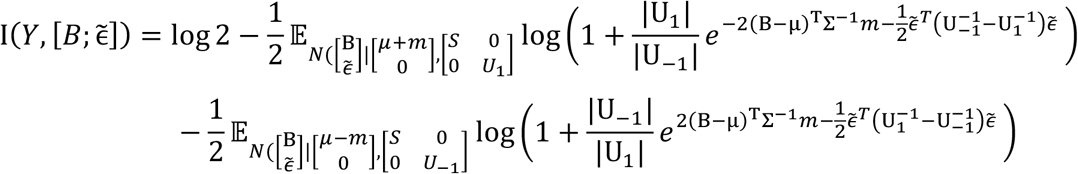

Now noting that log(1 + *e*^−X^) is a convex function, and that the expectations over B and 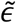 can be separated as they are independent, we can apply Jensen’s inequality to the expectation over 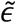 terms to obtain:

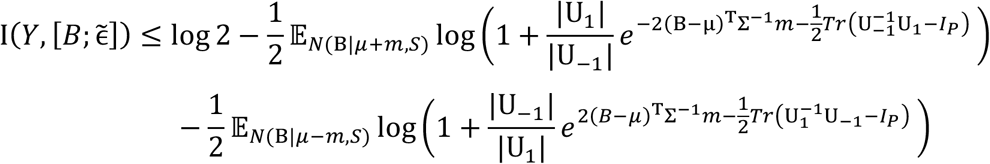

Let us now define a generalisation of the previously defined function *f*(*α*) to the following:

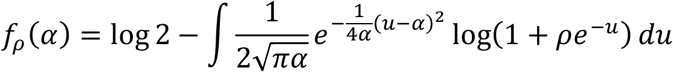

This allows us to write the information content upper bound as:

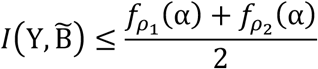

where *α* is the same term specified in Appendix A, and the new terms 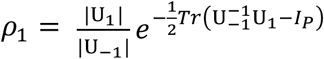 and 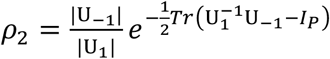.

### G. Upper bound for narrowband information content with induced effects

Let us now model the narrowband signal with stimulus dependent induced effects as follows:

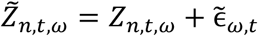

Where the new residual terms have distribution 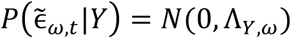. Note this is the same form as the model given in Appendix F, by substituting:

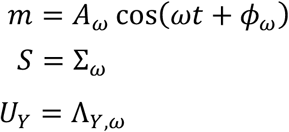

From Appendix F, we deduce the following:

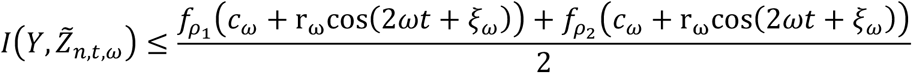

Where 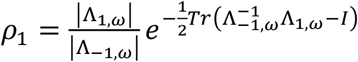 and 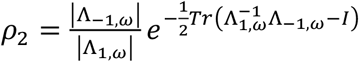 are both constant with respect to time, and *c*_*ω*_, *r*_*ω*_ and *ξ*_*ω*_ are the same terms specified in Appendix C. Thus, the narrowband information content associated with condition-dependent evoked *and* induced effects has an upper bound which is a sinusoidal function translated to double the original narrowband signal frequency (i.e. a slightly modified function of the same dynamics previously characterised for the case where only evoked effects are stimulus-dependent).

### H. Upper bound for broadband information content with induced effects

Let us now model the broadband signal with stimulus dependent induced effects as follows:

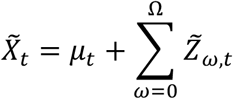

This is equivalent to the model of Appendix F by substituting:

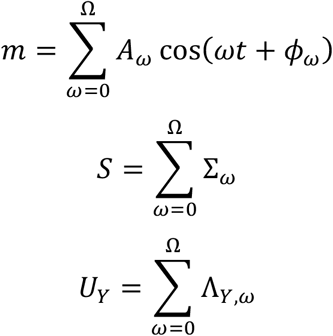

We therefore deduce that:

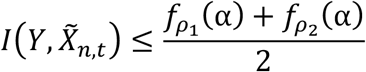

Where 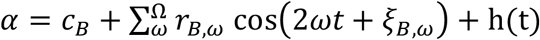, (i.e. the same as in the case of Appendix D where only evoked effects were modelled), *c*_*B*_, *r*_*B,ω*_, *ξ*_*B,ω*_ and h(t) are all as specified in Appendix D, and the new terms 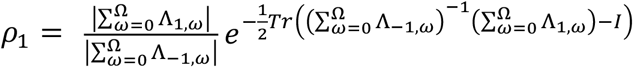 and 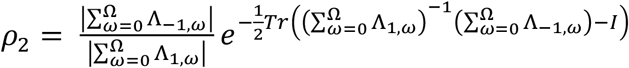 are both constant with respect to time.

## Acknowledgements

This research was funded by the Wellcome Trust (106183/Z/14/Z, 215573/Z/19/Z), the New Therapeutics in Alzheimer’s Diseases (NTAD) study supported by UK MRC and the Dementia Platform UK (RG94383/RG89702) and the EU-project euSNN (MSCA-ITN H2020-860563), and supported by the NIHR Oxford Health Biomedical Research Centre. The Wellcome Centre for Integrative Neuroimaging is supported by core funding from the Wellcome Trust (203139/Z/16/Z). DV is supported by a Novo Nordisk Emerging Investigator Award (NNF19OC-0054895) and by the European Research Council (ERC-StG-2019-850404). For the purpose of open access, the author has applied a CC BY public copyright licence to any Author Accepted Manuscript version arising from this submission.

## Declaration

The authors have no interests to declare.

## Data and code availability

The data used in Results Section 4 is from a previously published work (Cichy et al., 2016); it is publicly available for download at http://userpage.fuberlin.de/rmcichy/fusion_project_page/main.html. The code to perform all the analysis and example simulations published in this paper is publicaly available at https://github.com/OHBA-analysis/RepresentationalDynamicsModelling.

## Ethics Statement

The data used in Results Section 4 is from a previously published work (Cichy et al., 2016); as established in the original publication, the study was conducted in accordance with the Declaration of Helsinki and approved by the local ethics committee (Institutional Review Board of the Massachusetts Institute of Technology).

