## Supplementary Info for "The relationship between frequency content and representational dynamics in the decoding of neurophysiological data"

### Supplementary Information

#### S1. Generalising results to other common decoding metrics

In our analysis we have derived the spectrum of the information content up to an arbitrary monotonic scaling denoted by the function $f$. As this function is monotonic, it plays a role similar to – to use a more common example – mapping from a linear to a log scale. Importantly, whilst we have focussed on mutual information as a theoretical metric, this itself is has a monotonic relationship with many measures that are more commonly reported in decoding studies, such as classification accuracy or distance from the hyperplane (these are usually considered appropriate proxy measures of the mutual information). Thus, the relationships we have characterised extend equivalently to analysis using these metrics by simply exchanging the function (see Table 2 and Figure S1).

Strictly speaking, our results do not tell us the exact Fourier spectrum of these metrics, only that of the space prior to applying the appropriate monotonic scaling function. Nonetheless we consider this to be as informative, for two reasons. Firstly, as all of these functions are monotonic, they do not alter the timing of successive peaks and troughs in the metrics, nor the time period between them. Thus the major features of the response dynamics are preserved, the waveforms themselves are just warped slightly from a perfect sinusoid. Secondly, as implied above, this in fact makes the interpretation broadly applicable to a range of different metrics in commonly used in decoding analyses, as the same understanding applies regardless of which metric one chooses – each one just applies a slightly different warping to the sinusoidal component waveforms. This warping is characterised in Figure S1.

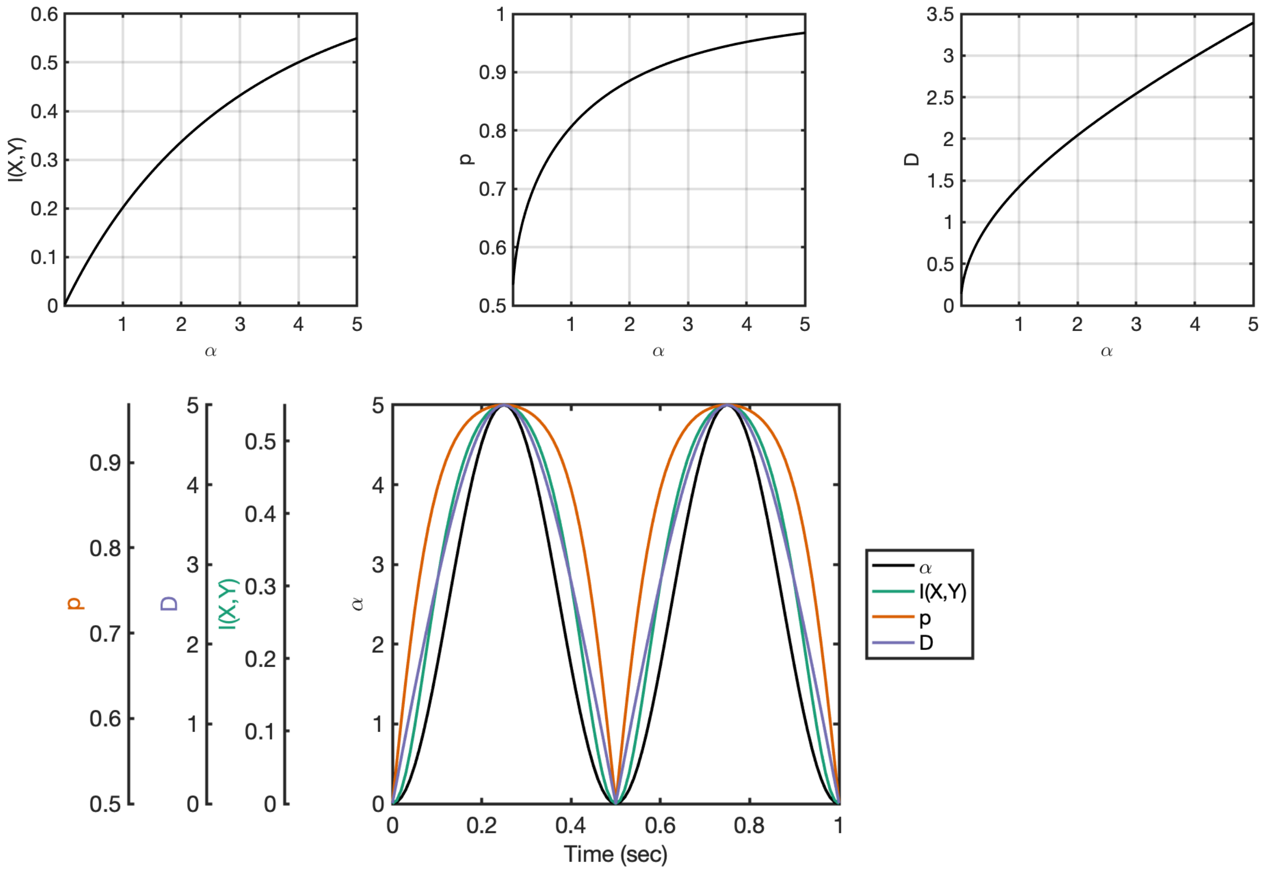


**Figure S1**: commonly reported decoding accuracy metrics are monotonic functions of the quantities modelled in Equation 3. Top panel: for some value $\alpha$, the scaling function $I(X, y) = f(\alpha)$ referred to in Equation 3. This same value has a one-to-one relationship with more common decoding measures such as classification accuracy (above centre) or distance from the hyperplane (above right). Lower panel: the relationships between these quantities can be further understood by plotting how an oscillation in $\alpha$ space maps into the mutual information space, the classification accuracy space or the hyperplane distance space – all are oscillatory with the same fundamental period, but with warping of the oscillatory shape between peaks and troughs (note the different scales on left for the different metrics; all scales are linear and scaled to match the extrema of the sinusoid in each space).

**S2. <u>Effect of window sizes on complex spectrum decoding</u>**

Complex spectrum decoding, when used with sliding window methods such as STFTs as we have done, is naturally subject to the trade-offs in time and frequency resolution that such techniques impose. This affects some of the interpretation argument that we have made. Specifically, in Results Section 3 we simulate non stationary and non-sinusoidal examples and show some of the benefits of complex spectrum decoding. In Figure 5 Example 1 we show complex spectrum decoding resolves the frequencies at which information appears in the signal as a function of time within epoch; in Example 2 we show that we can still identify two distinct activations that are not characterised parsimoniously by sinusoids. The main figure used a window size of 100msec.

To characterise the effect of window sizes on this interpretation, we ran the same analysis using window sizes of either 50msec or 250msec. Shorter windows would be expected to have poorer frequency resolution especially in lower frequencies, whilst larger windows would be expected to display poorer time resolution. Figure S2 confirms this is the case. The results for the shorter window are in this case not much affected; the results for the larger window (which is wider than the period between activations in Example 2) no longer allow disambiguation of the two different activations, which are now blurred together by the poorer time resolution imposed.

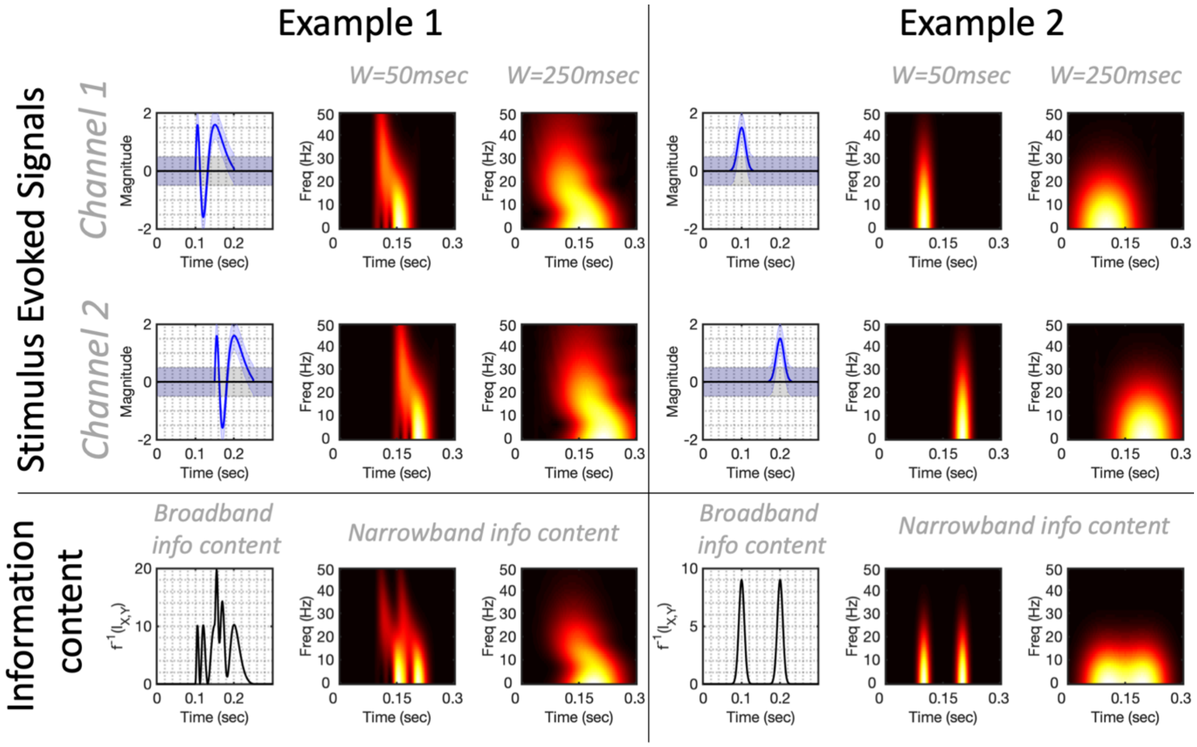


**<u>Figure S2: Replicating the analysis of figure 5 for wider or shorter windows.</u>** Top panel characterises the stimulus evoked signals which are the same as those presented in Figure 5; the time-frequency plots for these signals uses different windows demonstrating the classic time-frequency trade-off that window sizes impose for these techniques. Lower panel demonstrates the effects of window sizes on the associated information content evaluation by complex spectrum decoding.

**S3. <u>Derivation of higher moments of the distribution $Q(u)$</u>**

The moment generating function for the distribution $Q(u)$ is given as:

$$F(t) = \mathbb{E}_{Q(u)} e^{tu} = \frac{1}{C_1} M(t) G(t)$$

Where $M(t) = \mathbb{E}_{N(u|\alpha,2\alpha)} e^{tu}$, i.e. the moment generating function for the Gaussian distribution with mean $\alpha$ and variance $2\alpha$, and $G(t) = \mathbb{E}_{N(u|\alpha,2\alpha)} \log(1 + e^{-u-2\alpha t})$. From the properties of moment generating functions we can then obtain the $n$th moment of the distribution by the following:

$$\mathbb{E}_{Q(u)} u^n = \frac{d^n}{dt^n} F(t) \Big|_{t=0}$$

i.e. the $n$th derivative of $F(t)$ evaluated at $t = 0$. Denoting by $F^{(n)}(t)$ the $n$th derivative of $F(t)$, we have:

$$F^{(1)}(t) = \frac{1}{C_1}\left(M^{(1)}(t)G(t) + M(t)G^{(1)}(t)\right)$$

$$F^{(2)}(t) = \frac{1}{C_1}\left(M^{(2)}(t)G(t) + 2M^{(1)}(t)G^{(1)}(t) + M(t)G^{(2)}(t)\right)$$

$$F^{(3)}(t) = \frac{1}{C_1}\left(M^{(3)}(t)G(t) + 3M^{(2)}(t)G^{(1)}(t) + 3M^{(1)}(t)G^{(2)}(t) + M(t)G^{(3)}(t)\right)$$

$$F^{(4)}(t) = \frac{1}{C_1}\Big(M^{(4)}(t)G(t) + 4M^{(3)}(t)G^{(1)}(t) + 6M^{(2)}(t)G^{(2)}(t) + 4M^{(1)}(t)G^{(3)}(t) + M(t)G^{(4)}(t)\Big)$$

The higher moments $M^{(n)}(t)\big|_{t=0}$ for the Gaussian distribution are well established. Given a mean of $\alpha$ and variance of $2\alpha$ these are given by:

$$M(0) = 1$$

$$M^{(1)}(0) = \alpha$$

$$M^{(2)}(0) = \alpha^2 + 2\alpha$$

$$M^{(3)}(0) = \alpha^3 + 6\alpha^2$$

$$M^{(4)}(0) = \alpha^4 + 12\alpha^3 + 12\alpha^2$$

It remains then to evaluate the higher derivatives of the function $G(t)$. For simplicity of notation, we shall write the logistic sigmoid function as $\sigma(x) = \frac{1}{1+e^{-x}}$, and note the following key properties:

$$\sigma(x) = 1 - \sigma(-x)$$

$$\frac{d}{dx}\sigma(f(x)) = f'(x)\sigma\big(f(x)\big)\Big(1 - \sigma\big(f(x)\big)\Big)$$

Returning to the higher derivatives of $G(t)$, we have:

$$G^{(1)}(t) = \mathbb{E}_{N(u|\alpha,2\alpha)}\frac{\partial}{\partial t}\log(1 + e^{-u-2\alpha t})$$

$$= -2\alpha\mathbb{E}_{N(u|\alpha,2\alpha)}\sigma(-u - 2\alpha t)$$

Where we have made use of Leibniz's rule for differentiation under the integration sign. Evaluating at $t = 0$ we have:

$$G^{(1)}(0) = -2\alpha\mathbb{E}_{N(u|\alpha,2\alpha}(1 - \sigma(u))$$

We can then obtain the following:

$$G^{(2)}(t) = -2\alpha\mathbb{E}_{N(u|\alpha,2\alpha)}\frac{\partial}{\partial t}\sigma(-u - 2\alpha t)$$

$$= 4\alpha^2\mathbb{E}_{N(u|\alpha,2\alpha)}\sigma(u + 2\alpha t)\big(1 - \sigma(u + 2\alpha t)\big)$$

Such that

$$G^{(2)}(0) = 4\alpha^2\mathbb{E}_{N(u|\alpha,2\alpha)}\Big(\sigma(u)\big(1 - \sigma(u)\big)\Big)$$

Similarly:

$$G^{(3)}(t) = 4\alpha^2\mathbb{E}_{N(u|\alpha,2\alpha)}\frac{\partial}{\partial t}\big(\sigma(u + 2\alpha t) - \sigma^2(u + 2\alpha t)\big)$$

$$= 8\alpha^3\mathbb{E}_{N(u|\alpha,2\alpha)}\big(\sigma(u + 2\alpha t) - 3\sigma^2(u + 2\alpha t) + 2\sigma^3(u + 2\alpha t)\big)$$

Such that

$$G^{(3)}(0) = 8\alpha^3\mathbb{E}_{N(u|\alpha,2\alpha)}\big(\sigma(u) - 3\sigma^2(u) + 2\sigma^3(u)\big)$$

$$= 8\alpha^3\mathbb{E}_{N(u|\alpha,2\alpha)}(\sigma(u)(1 - \sigma(u))(1 - 2\sigma(u))$$

And:

$$G^{(4)}(t) = 8\alpha^3\mathbb{E}_{N(u|\alpha,2\alpha)}\frac{\partial}{\partial t}\big(\sigma(u + 2\alpha t) - 3\sigma^2(u + 2\alpha t) + 2\sigma^3(u + 2\alpha t)\big)$$

$$= 16\alpha^4\mathbb{E}_{N(u|\alpha,2\alpha)}\Big(\sigma(u + 2\alpha t)\big(1 - \sigma(u + 2\alpha t)\big)(1 - 2\sigma(u + 2\alpha t))^2$$

$$- 2\sigma^2(u + 2\alpha t)\big(1 - \sigma(u + 2\alpha t)\big)^2\Big)$$

Such that

$$G^{(4)}(0) = 16\alpha^4 \mathbb{E}_{N(u|\alpha,2\alpha)} \left(\sigma(u)\big(1-\sigma(u)\big)(1-2\sigma(u))^2 - 2\sigma^2(u)\big(1-\sigma(u)\big)^2\right)$$

Furthermore, note that $G(0) = \mathbb{E}_{N(u|\alpha,2\alpha)} \log(1+e^{-u}) = C_1$, i.e. the normalising constant as previously defined.

We can now evaluate the higher moments.

$$\mathbb{E}_{Q(u)} u^2 = F^{(2)}(t)\big|_{t=0}$$

$$= \frac{1}{C_1}\left(M^{(2)}(0)G(0) + 2M^{(1)}(0)G^{(1)}(0) + M(0)G^{(2)}(0)\right)$$

$$= \frac{1}{C_1}\left(C_1(\alpha^2+2\alpha) - 4\alpha^2 \mathbb{E}_{N(u|\alpha,2\alpha} 1-\sigma(u) + 4\alpha^2 \mathbb{E}_{N(u|\alpha,2\alpha)}\left(\sigma(u)\big(1-\sigma(u)\big)\right)\right)$$

$$= \alpha^2 + 2\alpha - \frac{4\alpha^2}{C_1}\mathbb{E}_{N(u|\alpha,2\alpha)}\big(1-\sigma(u)\big)^2$$

Similarly, we can obtain:

$$\mathbb{E}_{Q(u)} u^4 = F^{(4)}(t)\big|_{t=0}$$

$$= \frac{1}{C_1}\Big(M^{(4)}(0)G(0) + 4M^{(3)}(0)G^{(1)}(0) + 6M^{(2)}(0)G^{(2)}(0) + 4M^{(1)}(0)G^{(3)}(0)$$

$$+ M(0)G^{(4)}(0)\Big)$$

Evaluating and simplifying each term:

$$M^{(4)}(0)G(0) = C_1(\alpha^4 + 12\alpha^3 + 12\alpha^2)$$

$$4M^{(3)}(0)G^{(1)}(0) = 4(\alpha^3+6\alpha^2)\left(-2\alpha\mathbb{E}_{N(u|\alpha,2\alpha}(1-\sigma(u))\right)$$

$$= (-8\alpha^4 - 48\alpha^3)\mathbb{E}_{N(u|\alpha,2\alpha}(1-\sigma(u))$$

$$6M^{(2)}(0)G^{(2)}(0) = 6(\alpha^2+2\alpha)\left(4\alpha^2\mathbb{E}_{N(u|\alpha,2\alpha)}\left(\sigma(u)\big(1-\sigma(u)\big)\right)\right)$$

$$= (24\alpha^4 + 48\alpha^3)\mathbb{E}_{N(u|\alpha,2\alpha}\left(\sigma(u)\big(1-\sigma(u)\big)\right)$$

$$4M^{(1)}(0)G^{(3)}(0) = 4(\alpha)\left(8\alpha^3\mathbb{E}_{N(u|\alpha,2\alpha)}(\sigma(u)(1-\sigma(u))(1-2\sigma(u))\right)$$

$$= 32\alpha^4\mathbb{E}_{N(u|\alpha,2\alpha)}(\sigma(u)(1-\sigma(u))(1-2\sigma(u))$$

$$M(0)G^{(4)}(0) = 16\alpha^4\mathbb{E}_{N(u|\alpha,2\alpha)}\left(\sigma(u)\big(1-\sigma(u)\big)(1-2\sigma(u))^2 - 2\sigma^2(u)\big(1-\sigma(u)\big)^2\right)$$

Adding all above terms and factorising, we have:

$$\mathbb{E}_{Q(u)}u^4 = \mathbb{E}_{N(u|\alpha,2\alpha)}\Bigg(\alpha^4 + 12\alpha^3 + 12\alpha^2 + \frac{(1-\sigma(u))}{C_1}\big(8\alpha^4(\sigma(u)-1)(2\sigma(u)-1)(2\sigma(u)-1) + 48\alpha^3(6\sigma(u)-1)\big)\Bigg)$$

$$\mathbb{E}_{Q(u)}u^4 = \alpha^4 + 12\alpha^3 + 12\alpha^2 - \frac{8\alpha^3}{C_1}\mathbb{E}_{N(u|\alpha,2\alpha)}\left(\big(1-\sigma(u)\big)^2(\alpha(2\sigma(u)-1)^2 + 6)\right)$$

These are the two expressions for the higher moments $\mathbb{E}_{Q(u)}u^4$ and $\mathbb{E}_{Q(u)}u^2$ used in the proof of Appendix B.
